# Comparative phosphorylation map of Dishevelled3 (DVL3)

**DOI:** 10.1101/621896

**Authors:** Kateřina Hanáková, Ondřej Bernatík, Petra Ovesná, Marek Kravec, Miroslav Micka, Matěj Rádsetoulal, David Potěšil, Lukáš Čajánek, Zbyněk Zdráhal, Vítězslav Bryja

**Affiliations:** Central European Institute of Technology, Masaryk University, Brno, Czech Republic; National Centre for Biomolecular Research, Faculty of Science, Masaryk University, Brno, Czech Republic; Department of Experimental Biology, Faculty of Science, Masaryk University, Brno, Czech Republic; Department of Histology and Embryology, Faculty of Medicine, Masaryk University, Brno, Czech Republic; Institute of Biostatistics and Analyses, Faculty of Medicine, Masaryk University, Brno, Czech Republic; Department of Cytokinetics, Institute of Biophysics, Academy of Sciences of the Czech Republic, Brno, Czech Republic

**Author notes:** equal contribution (Kateřina Hanáková, Ondřej Bernatík). Corresponding authors at: Assoc. Prof. Vítězslav Bryja, PhD., Department of Experimental Biology, Faculty of Science, Masaryk University, Kotlářská 2, 611 37 Brno, Czech Republic; Assoc. Prof. RNDr. ZbyněkZdráhal, Dr. Central European Institute of Technology Masaryk University, Kamenice 5, 625 00 Brno, Czech Republic.

**Keywords:** Dishevelled, DVL3, phosphorylation, kinase, mass spectrometry, TTBK2, Wnt

## Abstract

In the presented study we analyze phosphorylation of human Dishevelled 3 (DVL3) induced by its previously reported (CK1ε, NEK2, PLK1, CK2α, RIPK4, PKCδ) and the newly identified (TTBK2, Aurora A) kinases. DVL3 contains 131 Ser/Thr whose phosphorylation generates complex barcodes underlying diverse DVL3 functions in Wnt pathways and other processes. We use quantitative mass spectrometry and via several complementary pipelines calculate site occupancies and quantify phosphorylation of >80 phosphorylated residues. In order to visualize the complex phosphorylation patterns, we design a novel visualization diagram, phosphoplot. Finally, we compare the individual sample and data processing approaches, identify their strengths and weaknesses. Subsequently, we verified a set of anti-phospho-DVL antibodies and were able to successfully confirm induction for several of the phosphorylation sites. From the biological point of view, our data represent an important reference point and a toolbox for further analysis of DVL functions and phosphorylation events that control them.

**One sentence summary:** The study provides a comprehensive comparison of the phosphorylation of DVL3 induced by eight Ser/Thr kinases and identifies phosphorylation signatures associated with individual kinases.

## INTRODUCTION

Wnt signaling pathway has been linked to an etiology of multiple developmental defects, inherited diseases and many types of cancer *(1)*. Wnt pathways can be divided into several main “branches”. The best studied (canonical) pathway depends on β-catenin and members of the LEF/TCF (lymphoid enhancer-binding factor/T-cell factor) family of transcriptional factors. In the absence of Wnt ligand, the intracellular level of free β-catenin is constantly low due to the activity of a degradation complex including adenomatous polypolis coli, Axin and glycogen synthase kinase (GSK) 3β. Upon Wnt binding destruction complex gets inhibited and allows accumulation and nuclear activity of β-catenin. However, Wnts can activate also the other, so called non-canonical Wnt pathways, which are β-catenin-independent and biochemically distinct from canonical Wnt signaling. There is strong evidence that several such pathways exist (for complete overview see *(2)*).

All known Wnt-induced pathways are transduced via seven transmembrane receptors Frizzleds *(3)* and intracellular protein Dishevelled (Dsh in *Drosophila*, DVL1-3 in human). DVL proteins consist of three structured domains: N-terminally located DIX domain, centrally located PDZ domain and C-terminally located DEP domain. Individual domains are linked by the intrinsically disordered regions and extended by approximately 200 amino acid long, also intrinsically disordered C-terminus.

There is a general agreement based on the genetic experiments that DVL plays a crucial role as a signaling hub in both Wnt/β-catenin as well as non-canonical Wnt pathways *(4)*. Not only that, DVL has been reported to have multiple other functions – such as docking of basal body *(5, 6)* and function and maintenance of primary cilia *(7)*, cytokinesis *(8)* or positioning of the mitotic spindle *(9)*, all possibly linked to the function of DVL in the regulation of centrosomal cycle *(10)*.

Intriguingly, despite the well documented role of DVL in the Wnt signaling and the growing evidence for its participation in additional cellular processes, the molecular mechanisms that define how DVL acts in the cell are almost unknown. However, it is believed that post-translational modifications, particularly phosphorylation, can represent a key component of such regulatory mechanism. DVL proteins are very rich in serine (Ser) and threonine (Thr); for example, DVL3 contains 131 Ser/Thr, which can be potentially phosphorylated. Well described consequence of the activation of both Wnt/β-catenin as well as non-canonical Wnt pathway is phosphorylation of DVL by the Wnt-induced <u>C</u>asein <u>k</u>inase 1 □ (CK1□) *(11–14)*. In addition to CK1ε, multiple other kinases have been reported to phosphorylate DVL in different contexts. For example, CK2α in both Wnt signaling pathways *(15–17)*, Polo-like kinase (PLK) 1 in the control of mitotic spindle *(9)*, Nima-related kinase (NEK) 2 in the centrosome *(10, 18)*, protein kinase C (PKC) δ in non-canonical Wnt signaling *(19)*, and receptor-interacting protein kinase (RIPK) 4 in the Wnt/β-catenin signaling *(20)*. Recent work *(10, 21, 22)* has identified using mass spectrometry based methods more than 50 Ser/Thr of DVL that are indeed phosphorylated. However, functional significance of most individual phosphorylation sites in DVL remains unclear. With respect to current understanding of PTMs in the intrinsically disordered proteins, it is reasonable to speculate that physiological function is achieved rather by a combination (“barcode”) of individual phosphorylated sites than by the phosphorylation a single amino acid.

In this study we analyzed and compared phosphorylation of DVL3 induced by eight individual Ser/Thr kinases: these included previously reported DVL kinases (6 kinases listed in the previous paragraph), Aurora A that was reported with DVL in the same complex *(23)* and TTBK2, DVL kinase identified in this study. We have applied complementary proteomic approaches to describe in detail and in a quantitative manner how individual kinases modify DVL3. This phosphorylation barcoding has identified unique as well as common phosphorylation patterns and provided a reference point for the interpretation of existing as well as any future studies analyzing phosphorylation of DVL. Last but not least, our work provided an example of the universal pipelines for the phosphorylation analysis of proteins modified in a complex pattern at tens of residues.

## RESULTS

### Identification of TTBK2 as a kinase acting upstream of DVL

Our previous work identified interactions between DVL3, CEP164, and TTBK2 kinase *(24, 25)*. To examine possible regulation of DVL by TTBK2, we have tested whether this poorly characterized centrosomal kinase phosphorylates DVL3 and DVL2 in HEK293 cells. As shown in Fig. 1A, TTBK2 co-expression induced a prominent electrophoretic mobility shift of DVL3 and DVL2 in a kinase activity-dependent manner, suggesting that TTBK2 is capable to efficiently promote phosphorylation of DVL. TTBK2 has been described as a kinase that primarily resides on the mother centriole where it regulates ciliogenesis *(25–31)*. Given the previous reports on DVL localization to centrosome and the possible DVL implication in ciliogenesis *(7, 10, 32, 33)* we have tested whether overexpressed DVL has the capacity to displace TTBK2 from the centriole. As shown in Fig. 1B (left panel), we confirmed, in line with earlier reports, the localization of TTBK2 to the mother centriole, which is however not affected by overexpression of DVL3 (Fig. 1B, right).

**Figure 1.**
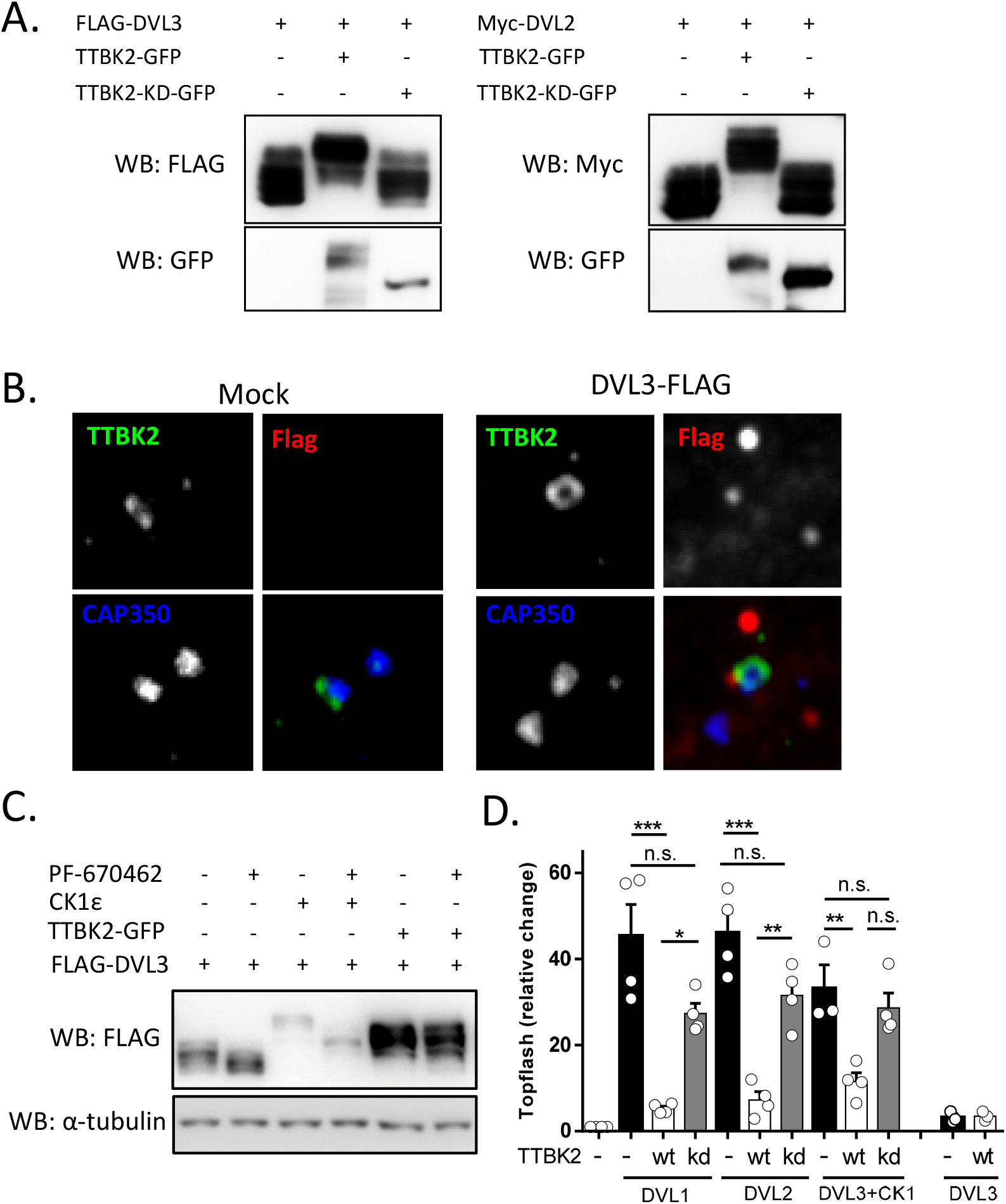
Identification of TTBK2 as a novel DVL kinase. A: HEK293 cells were transfected with FLAG-DVL3 and Myc-DVL2 plasmids with wild type (wt) or kinase dead (KD) TTBK2-GFP. Active TTBK2 promoted phosphorylation-dependent mobility shift of DVL3 on Western blotting. B: Endogenous TTBK2 (green) localized into distal appendages of the mother centriole in hTERT-RPE1 cells (left). Overexpression of FLAG-DVL3 (stained in red) was not able to displace TTBK2 from the centriole (right). Centrioles were stained with CAP350 (blue). C: HEK293 cells were transfected with indicated plasmids, treated with CK1ε inhibitor PF-670462 (10 μM) and subsequently analyzed by Western blotting. TTBK2-induced electrophoretic mobility shift of DVL3 was not diminished upon CK1ε inhibition unlike the mobility shift induced by CK1ε. D: HEK293 cells were transfected with indicated plasmids and by the TopFLASH reporter system. Luminescence in the cell lysates was measured 24 h after transfection. Mean, SD and individual data points are indicated. Statistical differences were tested by One-way ANOVA and Tukey’s post test (* p<0.05, ** p<0.01, *** p<0.001).

Mobility-shift of DVL2 and DVL3 induced by TTBK2 (Fig. 1A) can be in principle a consequence of direct phosphorylation of DVL by TTBK2 or a consequence of activation of other DVL kinase by TTBK2. In the second scenario the most established DVL kinase – CK1ε – represents the most obvious candidate target of TTBK2. To address whether TTBK2 acts directly on DVL or rather acts as the activator of CK1ε we treated cells with the CK1 inhibitor PF-670462 and subsequently analyzed the electrophoretic mobility shift of DVL3. As shown in Fig. 1C, PF-670462 efficiently reduced phosphorylation induced by CK1ε but not by TTBK2, hence demonstrating that TTBK2-induced phosphorylation of DVL does not require activity of CK1ε.

The best-defined role of DVL is the positive regulation of the Wnt/β-catenin pathway. In order to address if TTBK2 modulates this DVL function we have analyzed the ability of TTBK2 to promote or inhibit DVL-induced TCF/LEF-dependent luciferase reporter (TopFlash) in HEK293 cells. Interestingly, TTBK2 could in a kinase activity-dependent manner efficiently inhibit DVL1- and DVL2-induced TopFlash activation and could block positive effects of CK1ε on DVL3-induced TopFlash (Fig. 1D). DVL3 alone induces TopFlash very poorly (Fig. 1D, right). In summary, we describe robust phosphorylation of DVL mediated by TTBK2 that is associated with the decreased capacity of DVL to act in the Wnt/β-catenin pathway.

### Design and validation of DVL kinase panel

TTBK2 represents an addition to the ever-growing list of DVL kinases. Currently, multiple kinases from diverse kinase families have been reported to phosphorylate at least one of DVL isoforms. From the fragmented published results, it is not possible to find out what are the unique/general and constitutive/induced phosphorylation events and patterns associated with various functions of DVL in connection with individual kinases. This has inspired us to perform a direct comparison of DVL phosphorylation by individual kinases. We have chosen human HEK293 cells, a common model for the analysis of Wnt signaling, and human DVL3 as a representative DVL protein. Previously reported DVL kinases – CK1ε *(11)*, CK2α *(15)*, PLK1 *(9)*, NEK2 *(10, 18)*, PKCδ *(19)*, RIPK4 *(20)* and a newly identified TTBK2 were added to the panel. Finally, we have included also a mitotic kinase Aurora A that was reported to act in the same complex with DVL *(23)*. These Ser/Thr kinases represent very diverse members of the protein kinase family as visualized on the kinome tree (Fig. 2A).

**Figure 2.**
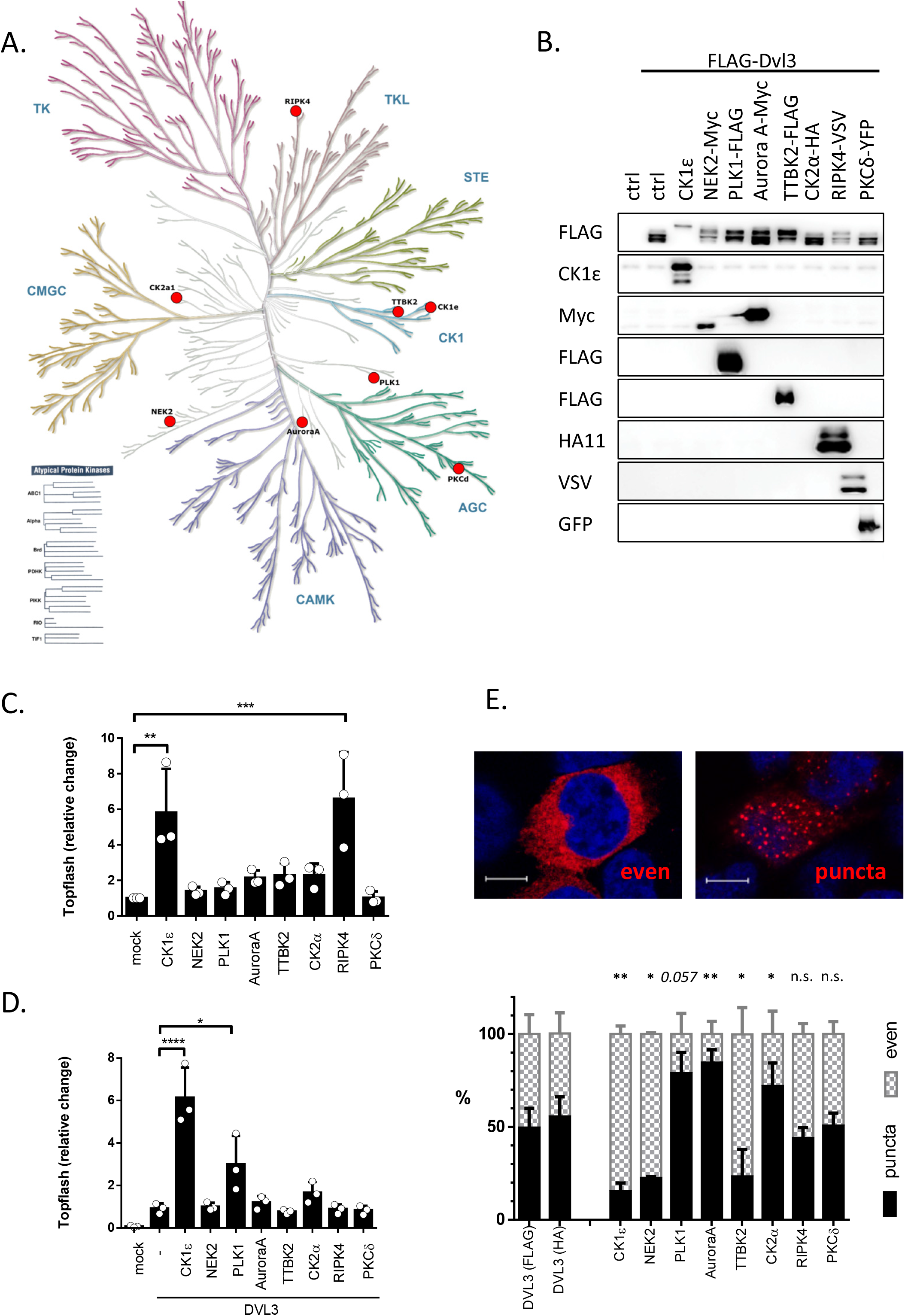
Validation of the panel of DVL3 kinases. A: Visualization of the kinases used in this study in the phylogenetic kinome tree (http://www.kinhub.org/kinmap/). The individual kinases are representatives of distant kinase groups except for CK1ε and TTBK2 that are members of CK1 superfamily. B. HEK293 cells were transfected by plasmids encoding FLAG-DVL3 and the indicated kinase. Ability of individual kinases to promote DVL3 phosphorylation detected as the electrophoretic mobility shift on WB was assayed. Alpha-tubulin was used as a loading control. C, D: HEK293 cells were transfected by indicated plasmids together with TopFlash and Renilla reporter plasmids. The ability of individual kinases to induce activation of Wnt/β-catenin pathway either alone (C) or in combination with DVL3 (D) was analyzed. Mean, SD and individual data points are indicated. Statistical differences were tested by One-way ANOVA and Tukey’s post test (* p<0.05, ** p<0.01, *** p<0.001, **** p<0.0001) E: HEK293 cells were transfected in the indicated combinations and the subcellular localization of DVL3 was assessed by immunocytochemistry. DVL3 was localized in two typical patterns – either in cytoplasmic puncta or evenly dispersed in the cytoplasm (upper panel). Scale bar, 7.5 μm. The effects of individual kinases on DVL3 localization is shown in the bottom panel (HA-DVL3 was used for PLK1 and TTBK2, FLAG-DVL3 for the rest of kinases). The data represent mean + SD from three independent experiments (N=3× 200 cells). Statistical significance was confirmed by the comparison of the corresponding control (DVL3-FLAG or DVL3-HA without kinase) and DVL3 with individual kinases by One-way ANOVA and Tukey’s post test (* p<0.05, ** p<0.01, n.s. - not significant).

To set up a starting point for the subsequent proteomic analyses and to validate the performance of selected kinases towards DVL, we have compared individual kinases in three assays: (i) capacity to induce phosphorylation-dependent mobility shift of DVL3, (ii) capacity to induce Wnt/β-catenin-dependent transcription analyzed by the TopFlash reporter assay, and (iii) capacity to change the subcellular localization of DVL3.

In the first assay, most kinases – except for CK2α and PKCδ – could trigger electrophoretic mobility shift of DVL3 when co-expressed with DVL3. In the TopFlash reporter assay that reflects the activation of downstream events in the Wnt/β-catenin pathway, only CK1ε and RIPK4 were capable to induce activation in the absence of exogenous DVL3 (Fig. 2C). In the presence of exogenous DVL3, only CK1ε, and to a lesser extent PLK1 (non-significant trend was observed also for CK2α), could synergize with the co-expressed DVL3 to promote reporter activation (Fig. 2D). In the third assay we have analyzed the capacity of individual kinases to alter the subcellular localization of DVL3. Overexpressed DVL3, as well as other DVL proteins, is most commonly localized in the so called “DVL punctae” protein assemblies kept together via polymerization of DVL DIX domains *(34)*. These assemblies are dynamic and depending on the stimuli they can be more compact (visible as puncta) or dissolved, resulting in the “even” distribution of DVL3 – see Fig. 2E, upper panel. As shown in Fig. 2E, bottom panel, three kinases – CK1ε, NEK2 and TTBK2 – were capable to promote even localization of DVL3 whereas Aurora A and CK2α (and to some extent also PLK1) lead to a more punctate phenotype.

This initial analysis showed that we can largely reproduce the reported effects of selected kinases: the capacity of CK1ε and RIPK4 to activate TopFlash reporter *(20, 35)*, the synergistic behavior of DVL3 and CK1ε in the TopFlash reporter *(17)*, and the capacity of CK1ε and NEK2 to induce even distribution of DVL3 *(10, 13)*. CK1ε stands unique among other kinases in its ability to efficiently induce Wnt/β-catenin downstream signaling in synergy with DVL3. Although, it has been proposed that RIPK4 activate Wnt/β-catenin also via phosphorylation of DVL *(20)* we have not observed any synergy with DVL3 which suggests that RIPK4 acts via other proteins in the Wnt/β-catenin pathway.

### Pipelines for the generation of the phosphorylation map of DVL3

Following validation of our experimental system (Fig. 2) we moved to the global analysis of the DVL3 phosphorylation events. In three independent experiments, FLAG-DVL3 was overexpressed in HEK293 cells, with or without the studied kinase, immunoprecipitated using anti-FLAG antibody, separated on SDS-PAGE and stained with Coomassie Brilliant Blue (see Supplementary Fig. 1 for the gels used in this study). Bands corresponding to DVL3 were cut out and after TrypChymo digestion 1/10 of the sample was analyzed directly (see pipeline #1 in Fig. 3A) and the remaining 9/10 of the peptide mixture was enriched for the phosphorylated peptides by TiO_2_ (see pipelines #2 and #3). All samples were subsequently analyzed by liquid chromatography coupled with mass spectrometry (LC-MS/MS).

**Figure 3.**
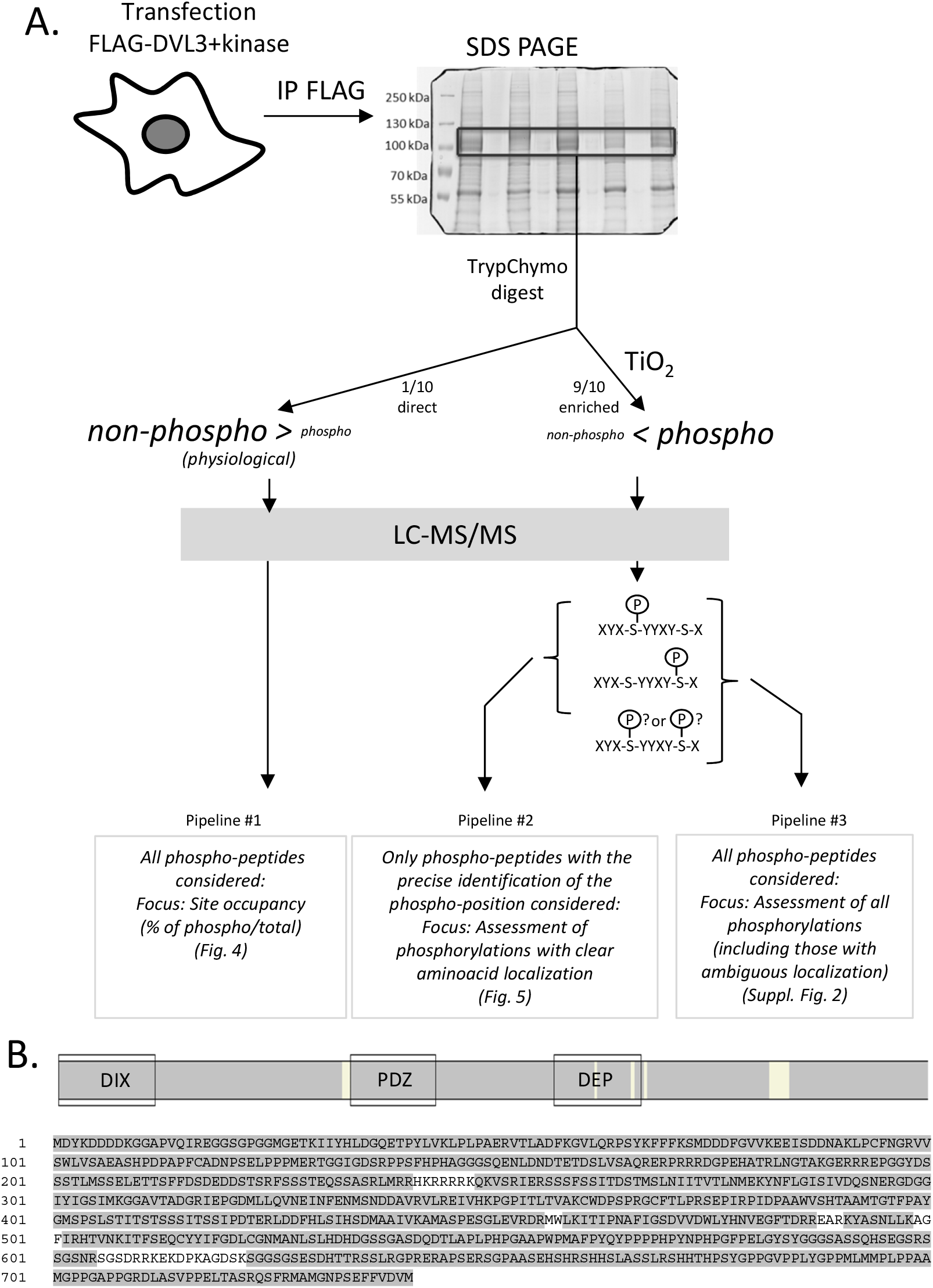
Experimental design. A: FLAG-DVL3 was overexpressed (with or without kinase) in HEK293 cells. After cell lysis DVL3 was immunoprecipitated using anti-FLAG antibody. Immunoprecipitates were separated on SDS-PAGE gel electrophoresis, stained with Coomassie brilliant blue and the 1D bands corresponding to DVL3 were excised, digested with trypsin and subsequently cleaved by chymotrypsin. In the pipeline #1, the aliquot (1/10) of concentrated sample was directly analyzed by LC-MS/MS in order to analyze site occupancy of the abundant phosphorylated sites. The rest of the sample was enriched for phosphorylated peptides using TiO_2_ and analyzed by LC-MS/MS to obtain detailed information of DVL3 phosphorylation status. Data from LC-MS/MS were searched, manually validated in Skyline software and further processed by two approaches. In the first approach (pipeline #2) only phosphorylated peptides with the clearly localized phosphorylated site (based on manual inspection of spectra) were considered. In the second approach (pipeline #3) we have considered all phosphorylated peptides that in some cases resulted in the formation of “clusters” of phosphorylated sites. B. The overall sequence coverage of DVL3 across all kinases and replicates. Regions of DVL3 covered by the peptides detected (Mascot score > 20) in any of the MS/MS analyses are highlighted in grey. For sequence coverage in individual samples see Suppl. Table 2.

Peptides with accurately characterized phosphorylated site(s) were identified alongside peptides where the phosphorylation localization was not reliably assigned, as is common to proteomic studies. Therefore, in the accurate position focused analysis (pipeline #2) only a subgroup of peptides with the clearly identified phosphorylated positions (based on manual inspection) was included. In the overall quantitative analysis (pipeline #3) all phosphorylated peptides were processed despite the positions of the phosphorylated residues were not always certain and as such are presented as the “phosphorylated clusters”. For the detailed information on the quantification in these two pipelines see Materials and Methods.

Combination of the above-mentioned approaches allowed us to cover more than 95% of the DVL3 sequence across all experiments (Fig. 3B); for sequence coverage in individual samples see Suppl. Table 2.

### Phosphorylation map of DVL3: site occupancy of phosphorylated sites

To assess the site occupancy, we utilized direct analysis of the samples without any enrichment (the experimental pipeline #1, Fig. 3A) which allowed us to detect phosphorylated and non-phosphorylated peptides corresponding to the same position(s). Subsequently we calculated the approximate occupancy of the selected phosphorylated sites, i.e. % of DVL3 molecules phosphorylated at the given Ser or Thr residue(s).

We calculated site occupancies as percentage ratio of phosphorylated peptide intensity (or sum of intensities if more than one) covering individual phosphosite or cluster to total intensity (phosphorylated peptide(s) + corresponding non-phosphorylated peptide(s)). Phosphorylated sites/clusters with the site occupancy > 5 % at least in one sample are plotted in Fig. 4. For three clusters (S232□S244, T608□S612, S622□S630) we have failed to detect matching non-phosphorylated peptides and as such these sites were not included in the site occupancy calculations.

**Figure 4.**
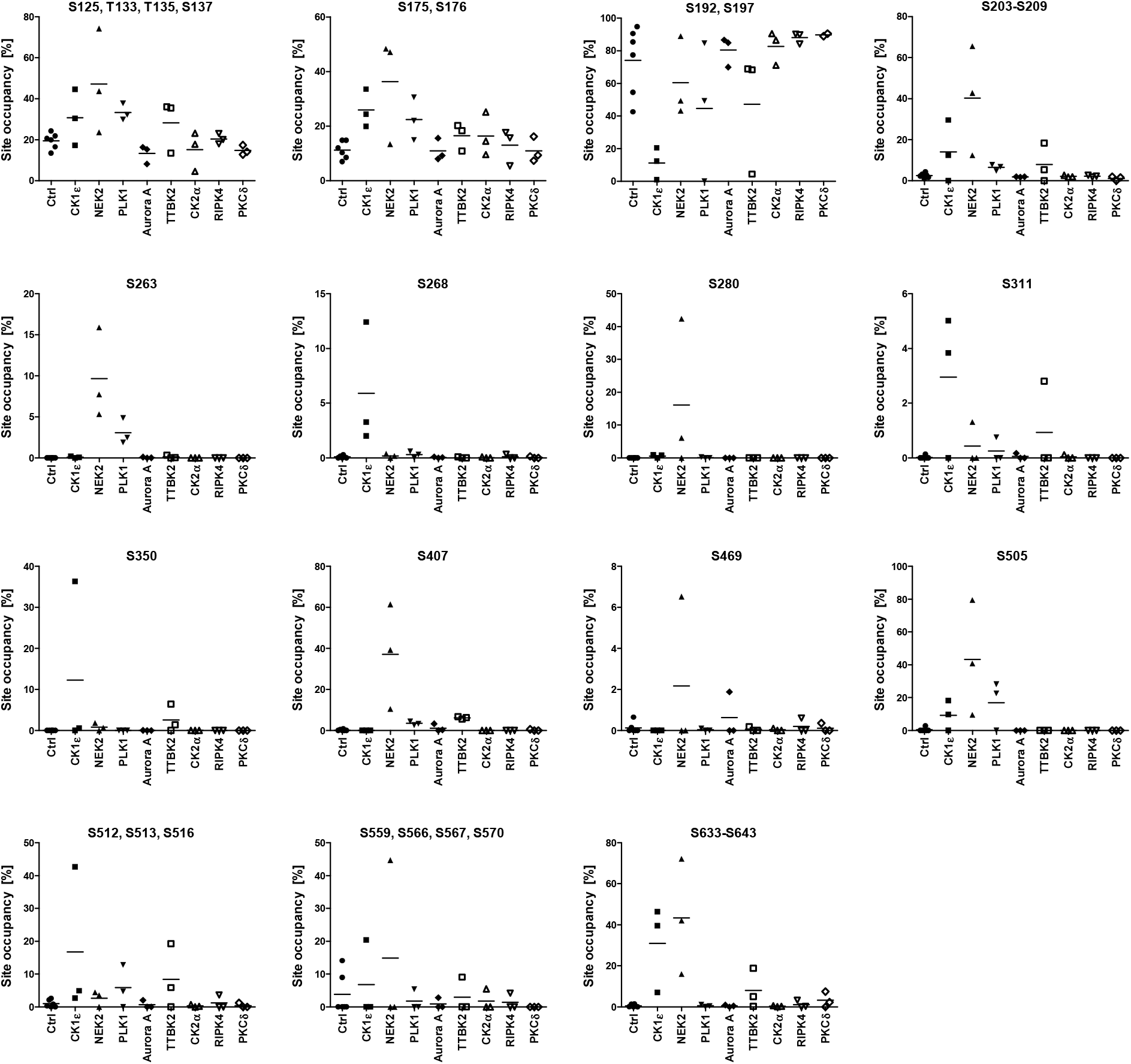
Site occupancy of the abundant phosphorylated sites. Site occupancy analysis was performed according to the pipeline #1 in Fig. 3A. Fifteen phosphorylated peptides or clusters that were phosphorylated in more than 5 % at least in one replicate are plotted. Graphs present individual data points from three biological replicates (two controls/biological sample) and the mean values (horizontal line).

We have observed an increase in the phosphorylated site occupancy (above 5%) after overexpression of at least one protein kinase in all but one *(14)* cases (Fig. 4). The notable exception was the phosphorylated cluster S192, S197, whose phosphorylation decreased in the presence of CK1ε and to a lesser extent in the presence of TTBK2, PLK1 and NEK2. Phosphorylation of three sites was induced selectively only by one kinase – this is the case of S280 and S407 induced only by NEK2, and S268 induced only by CK1ε. In the remaining 11 sites/clusters, two or three kinases were capable to trigger phosphorylation: CK1ε/TTBK2 at S350 and at the cluster S512, S513, S516; NEK2/PLK1 at S263; NEK2/Aurora A at S469 (in the single replicate); NEK2/PLK1/CK1ε at S505; NEK2/CK1ε/TTBK2 (and to some extent PLK1) at S311, and at the clusters S203□S209 and S633□S643. Other three clusters – cluster S125, T133, S135, S137, cluster S175, S176 and cluster S559, S566, S567, S570 showed relatively high occupancy in the control and were further phosphorylated by several kinases, including NEK2, CK1ε, TTBK2 and PLK1.

### Phosphorylation map of DVL3: detailed analysis after phospho-enrichment

In order to analyze phosphorylation of DVL3 in depth, we enriched the phosphorylated peptides by TiO2 enrichment. This approach, commonly utilized for detailed screening of protein phosphorylations, allows detection of less abundant phosphopeptides. The obtained data were processed in two ways (Fig. 3A). Primarily, we qualitatively and quantitatively assessed only phosphopeptides with the clearly localized (validated by manual inspection of MS/MS data) phosphorylation site(s) (the experimental pipeline #2, Fig. 3A). By this approach we have detected 88 unique phosphorylation sites in DVL3. Peptide intensities (Supplementary Table 1) were compared with the DVL3-only control dataset in order to express the relative increase/decrease in the phosphorylation of each phosphorylated site. Data from all three replicates presented as a relative change to control were summarized as a heat map in Fig. 5. Most phosphorylated sites with the biggest relative increase were identified after induction by NEK2, CK1ε and TTBK2.

**Figure 5.**
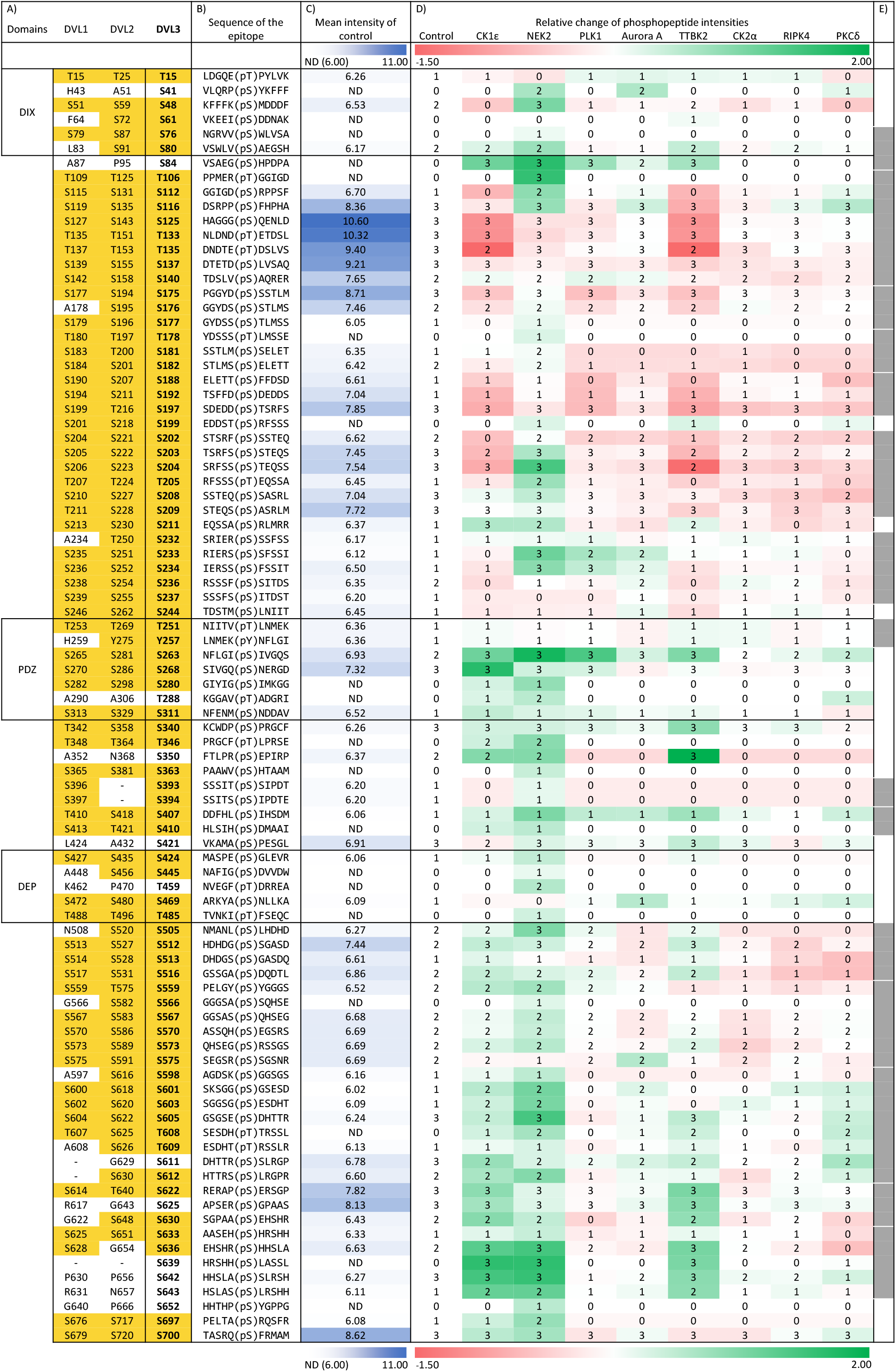
Phosphorylation map of DVL3. All identified phosphorylation sites obtained from the pipeline #2 are visualized as a heatmap. Color intensities reflect relative change in the site phosphorylation (red – decrease, green – increase). Following additional information is also provided: A) Corresponding sites in human DVL1 and DVL2. Positions conserved in DVL1 and/or DVL2 either as Ser or Thr are highlighted in yellow. Position of the structured domains (DIX, PDZ and DEP) is indicated. B) The sequence of the phosphorylated epitope. Five amino acids before and after the identified phosphorylated site are shown. C) Mean absolute intensities of the phosphorylation sites in the control (DVL3 without exogenous kinase; N=6) are expressed in the shades of blue. Numbers indicate decadic logarithm of the mean. ND indicated in white corresponds means “not detected”. All signals lower than 1×10^6^, corresponding to log value 6.0 were considered as not detected. D) Nine columns represent heat map of relative change of phosphorylated peptide intensities (in log10 scale) obtained for individual kinases (relative to control). Numbers in the heatmap fields (0, 1, 2, 3) indicate the number of experimental replicates with the positive identification of the given phosphorylated site. E) Grey boxes indicate the position of clusters of sites analyzed also according to the pipeline #3 in the Suppl. Fig. 2.

In order to assess impact of exclusion of phosphopeptides with ambiguous phosphosite localization on quantitative changes we have processed all phosphorylated peptides according to the pipeline #3 (Fig. 3A). Intensities of the phosphorylated clusters that were formed as the outcome of this approach, presented as a heat map in Supplementary Fig. 2, illustrated that intensities of the 15 clusters analyzed very well correlate with the intensities of individual phosphorylated sites presented in Fig. 5. Comparison of results in Fig. 5 and Suppl. Fig. 2, however, also identified cases where individual phosphorylated sites within the cluster displayed a distinct behavior. Namely, in the cluster T106□S140 the NEK2-induced phosphorylation of T106, S112 and S116 in the cluster was masked by high intensity constitutive phosphorylation of S125, T133 and S137. In the phosphorylated cluster S202□S209 that decreased in the presence of CK1ε and TTBK2 and increased for NEK2 we can map the decrease to three sites – S202, S203 and S204 – in case of CK1ε and TTBK2 whereas NEK2 induced predominantly S204. Similarly, for the cluster S598□S612 induced by NEK2, CK1ε, TTBK2 and PKCδ we mapped the activity of NEK2 predominantly to S601, S603 and S605 whereas CK1ε phosphorylated mainly S611 and S612.

The quantitative data presented in Fig. 5 (accurate positions) and Suppl. Fig. 2 (clusters) have been combined and analyzed in order to group both the individual phosphorylated sites/clusters and individual kinases. Unbiased cluster analysis is shown in Suppl. Fig. 3. It clearly clusters CK1ε and TTBK2, both from the CK1 family, with NEK2, whereas all the remaining kinases formed a second cluster. There are multiple residues that are phosphorylated only by NEK2, which clearly distinguishes it from CK1ε and TTBK2. Interestingly, CK1ε and TTBK2 behave very similarly and they can be best resolved by the phosphorylation at S268 induced only by CK1ε. This is in a very good agreement with the site occupancy analysis shown in Fig. 4.

### Phosphorylation map of DVL3: Analysis of the phosphorylated clusters

The phosphorylated clusters visualized in Suppl. Fig. 2 can in principle represent the mixtures of peptides phosphorylated at distinct sites or true multiphosphorylated motives with possible biological function. To get a better insight into this phenomenon, we analyzed in detail the peptides phosphorylated on 3 or more sites. We have found 9 peptide families fulfilling these requirements (Fig. 6). Out of these only one cluster – S112, S116, S125, T133, S135, S137 – was not induced by any of the kinases. Remaining 8 clusters are induced by one or several kinases – 3x only by NEK2, 2x mainly by CK1ε and TTBK2 and 3x by NEK2 and one or more additional kinases.

These phosphorylated clusters were detected only in the intrinsically disordered regions of DVL3. The phosphorylated motives are also surprisingly well conserved and with the exception of S622□630 and S636□S643 are found also in DVL1 and DVL2 (Fig. 6, right). This may suggest that the function of multiple phosphorylation of these motives in the regulation of DVL is also conserved.

### Comparison of individual pipelines and validation by the phospho-specific antibodies

In our study, we have used several sample and data processing pipelines (see Fig. 3). As the last step we have decided to compare individual pipelines (i) between each other and (ii) with several phosphorylation-specific antibodies raised against phosphorylated DVL3 peptides. We have probed the samples described in Fig. 2B with the antibodies against phosphorylated-Ser280-DVL3 (pS280) *(10)*, pS643 *(21)*, pS697 *(10)* and the newly generated anti-pS192. As we show in Fig. 7A-D that combines the Western blots and the MS/MS data from pipelines #1 and #2, the signal of phospho-specific antibodies partially matches the changes observed by mass spectrometry. Signal of pS192-DVL3 decreased after CK1ε, PLK1 and TTBK2 co-expression whereas pS280, pS643 and pS697 increased the most with NEK2. However, some differences were observed, especially for the lower intensity signals – namely, all three previously validated antibodies pS280, pS643 and pS697 do detect higher phosphorylation after Aurora A co-expression that was not detected as increased in these positions by LC-MS/MS. On the other hand – increase in pS643 convincingly demonstrated for TTBK2 by all MS/MS pipelines was missed by anti-pS643-DVL3 antibody.

**Figure 6.**
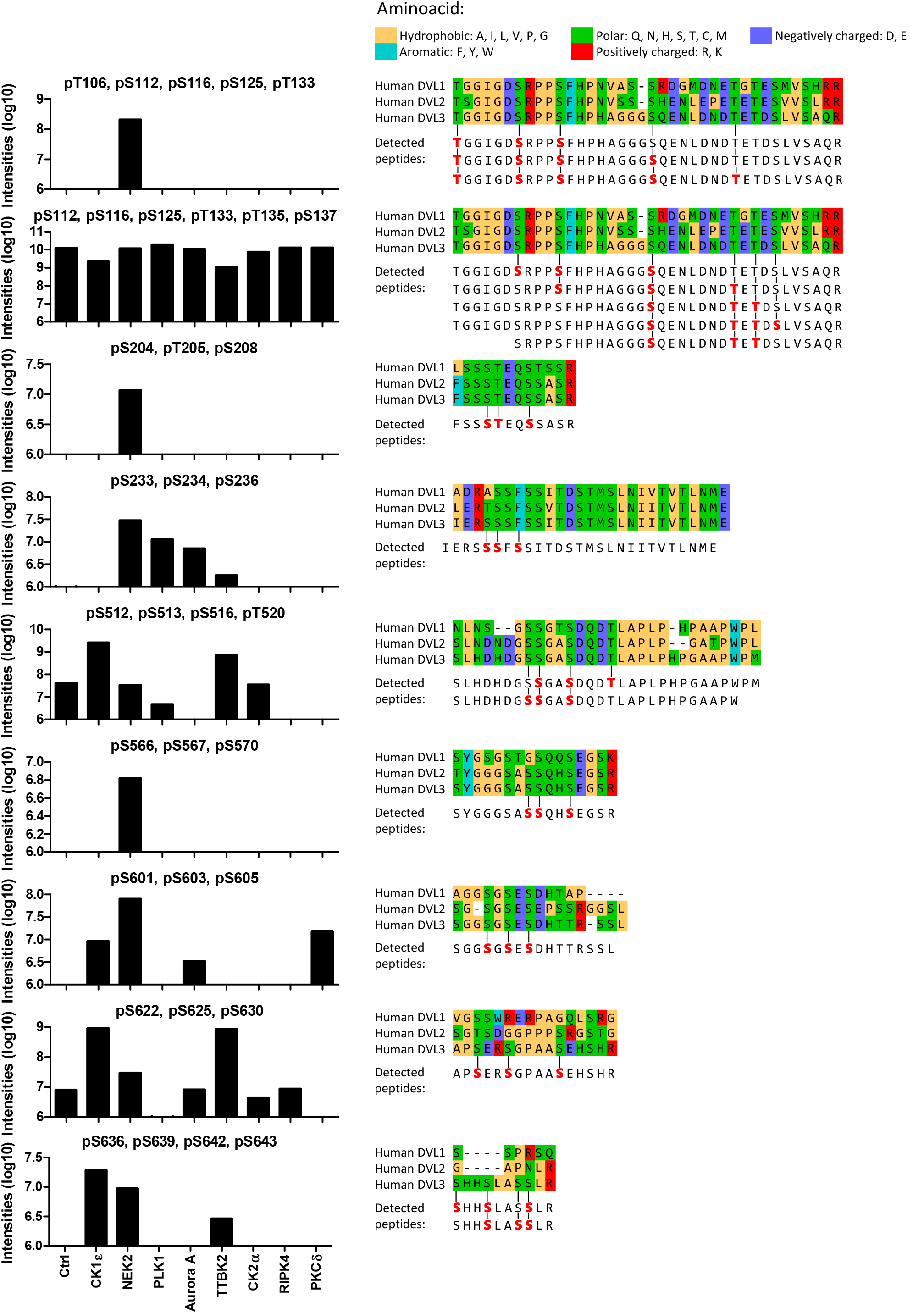
Clusters phosphorylated in more than three sites. The clusters of Ser/Thr where 3 or more phosphorylated sites in one peptide were analyzed (i.e. multiphosphorylated peptides). Graphs indicate total intensities of the multiphosphorylated peptides from all three replicates; signal intensities from pipeline #1 and #3 are merged. Multiple sequence alignment shows the evolutionary conservation of the motif among individual DVL isoforms. All combinations of individual multiphosphorylated peptides detected in the cluster are indicated.

### Phosphoplot – a tool for visualization of complex phosphorylation patterns

The heat map data visualization highlights the differences in the phosphorylation in comparison to control but does not provide the full and intuitive picture combining absolute intensities, their changes and position of individual phosphorylated peptides. To address this, we have designed a visualization diagram, that we refer to as a “phosphoplot”. In the phosphoplot all Ser/Thr from the primary sequence of the analyzed protein are shown. The intensities of phosphorylated peptides in the control and experimental conditions are plotted. As such phosphoplot combines information on absolute peptide intensities, experimental differences and positional information, including non-phosphorylated sites.

The phosphoplots of DVL3 for individual kinases based on data from pipeline #2 are shown in Fig. 8. The short inspection of individual phosphoplots identified regions of DVL3 that are devoid of phosphorylation, despite being Ser/Thr rich – such as T365□T392, and on the contrary the regions between the PDZ and DEP domain that are highly constitutively phosphorylated. It also clearly identifies the uniquely phosphorylated sites for each kinase.

## DISCUSSION

Our study provides the first comprehensive description of the phosphorylation of DVL3 by most of the described DVL Ser/Thr kinases. Given the high sequence conservation of DVL3 and other DVL proteins it is an important reference point for the interpretation of the published data as well as benchmark for forthcoming studies focused on the regulation of DVL function by kinases and phosphatases.

The parallel application of several pipelines for sample preparation and data analysis led us to conclusion that the occupancy of the phosphorylated site(s) provides a suitable approach for the identification of the phosphorylation events that could be biologically relevant. Site occupancy analysis clearly has the potential to detect phosphorylation events that could trigger biological response controlled by the kinase(s). Site occupancy analyses, that identified S268 as signature site for CK1ε, S280 as a signature site for NEK2 and S633□S643 cluster as a signature for CK1ε, NEK2 and TTBK2, serve as excellent examples. All these phosphorylation changes have been validated by additional, in every respect more demanding proteomic pipelines – compare Fig. 4 with Fig.5, Fig. 6 and Suppl. Fig. 3. Direct analysis of the sample by the site occupancy analysis should be thus considered as a valid approach in the biology-oriented studies with the aim to find candidate phosphorylation events responsible for the function associated with the given kinase in other Ser/Thr rich proteins with complex phosphorylation pattern.

The data collected by various approaches shown in Figs 4–7, provide a global view of DVL3 phosphorylation induced by kinases that were shown to control distinct functions of DVL. We can clearly observe distinct codes of phosphorylated Ser/Thr in the structured domains – most typically in the PDZ domain at the fully conserved S263, S268, S280 and S311, and to a lesser extent also in the DIX and DEP domains. On the other side, in the largely intrinsically disordered regions in between domains we have detected conserved (see Fig. 6) multiphosphorylated sequences with specific phosphorylation patterns. It is tempting to speculate that the multi phosphorylation in these motifs will have biochemical and/or biophysical consequences resulting in the regulation of DVL function. Most intrinsically disordered proteins or regions are very Ser/Thr rich because these aa (together with A, R, G, Q, P, E and K) are disorder-promoting. Intrinsically disordered proteins (such as DVL) are essential components of multiple signal transduction pathways *(36)* and a complex multiphosphorylation that we started to uncover in DVL can represent a shared and universal mechanism for the regulation of their function.

**Figure 7.**
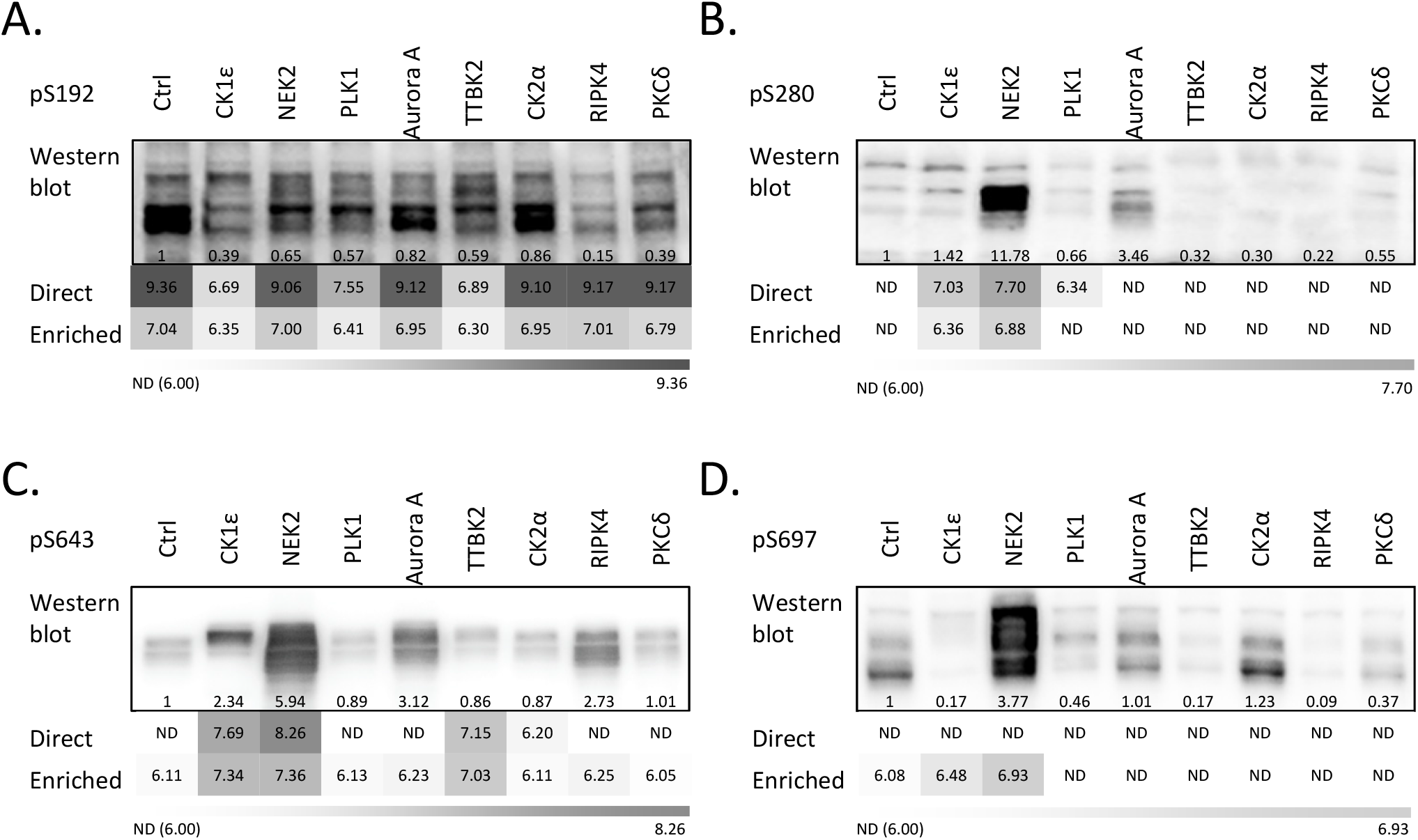
Comparison of individual methods and their validation by phospho-specific antibodies. A-D. Comparison of methods. Existing phosphorylation-specific antibodies were used to visualize the level of phosphorylation of DVL3 with our kinase panel. HEK293 cells were transfected by the indicated combination of plasmids and analyzed by Western Blotting. Reactivity of individual phosphoantibodies against DVL3 phosphorylated by individual kinases is shown. Western blots are quantified using ImageJ software as absolute values for peak areas of corresponding bands and the intensities were normalized to the control. Mean intensities of the phosphorylated peptides obtained by MS/MS via pipelines #1 and #2 are shown in the shades of grey. Numbers indicate decadic logarithm of the mean peptide intensity. “Not detected” (ND) indicated in white corresponds to the signals below 1×10^6^, i.e. 6.0. A. anti-pS192 (this study), B. anti-pS280, C. anti-pS643, D. anti-pS697.

An interesting phosphorylation pattern has been observed for two highly conserved regions in between DIX and DEP domain – corresponding to S112□S140 and S188□S197. These sites showed high level of basal phosphorylation in the absence of any exogenously coexpressed kinase. Interestingly, expression of CK1ε (and to a lesser extent TTBK2 and PLK1) dramatically reduced phosphorylation of these motifs. This suggests that there is a so far unidentified endogenous kinase that very efficiently phosphorylates these regions, and (ii) that the binding and/or phosphorylation by CK1ε and TTBK2 interferes with this process or perhaps promotes removal of these phospho-moieties by activation of specific phosphatase(s). So far only protein phosphatase 2A has been shown to have a positive function in the Wnt/β-catenin signaling upstream of DVL *(37, 38)*, which makes it an ideal candidate for such function.

Three kinases from our panel, CK1ε, NEK2 and TTBK2, were the only ones that phosphorylated cluster S630-S643. This capacity of CK1ε, NEK2 and TTBK2 correlated with the ability to trigger the even distribution of DVL3 in the cytoplasm, shown in this study as well as in the earlier reports *(10, 12, 13)*. This is line with the earlier work that showed the requirement of phosphorylated S630–S643 for the even distribution of DVL3 induced by CK1ε *(21)*. Phosphorylation-specific antibody that recognizes this cluster-(pS643) shows even cytoplasmic staining *(21)*, which further substantiates the importance of phosphorylated S630–S643 as a molecular marker of evenly distributed DVL3. On the other hand, these three kinases stood out as the most efficient DVL3 kinases when perceived from the global point of view. These kinases could very efficiently phosphorylate multiple Ser/Thr-rich clusters in the unstructured regions of DVL (see Fig. 5, 6 and 8). This opens the possibility that the even distribution of DVL3 is not only due to the phosphorylation at the specific Ser or Thr but rather reflects the consequence of massive phosphorylation of multiple residues accompanied by the change in the charge and protein biophysical properties.

**Figure 8.**
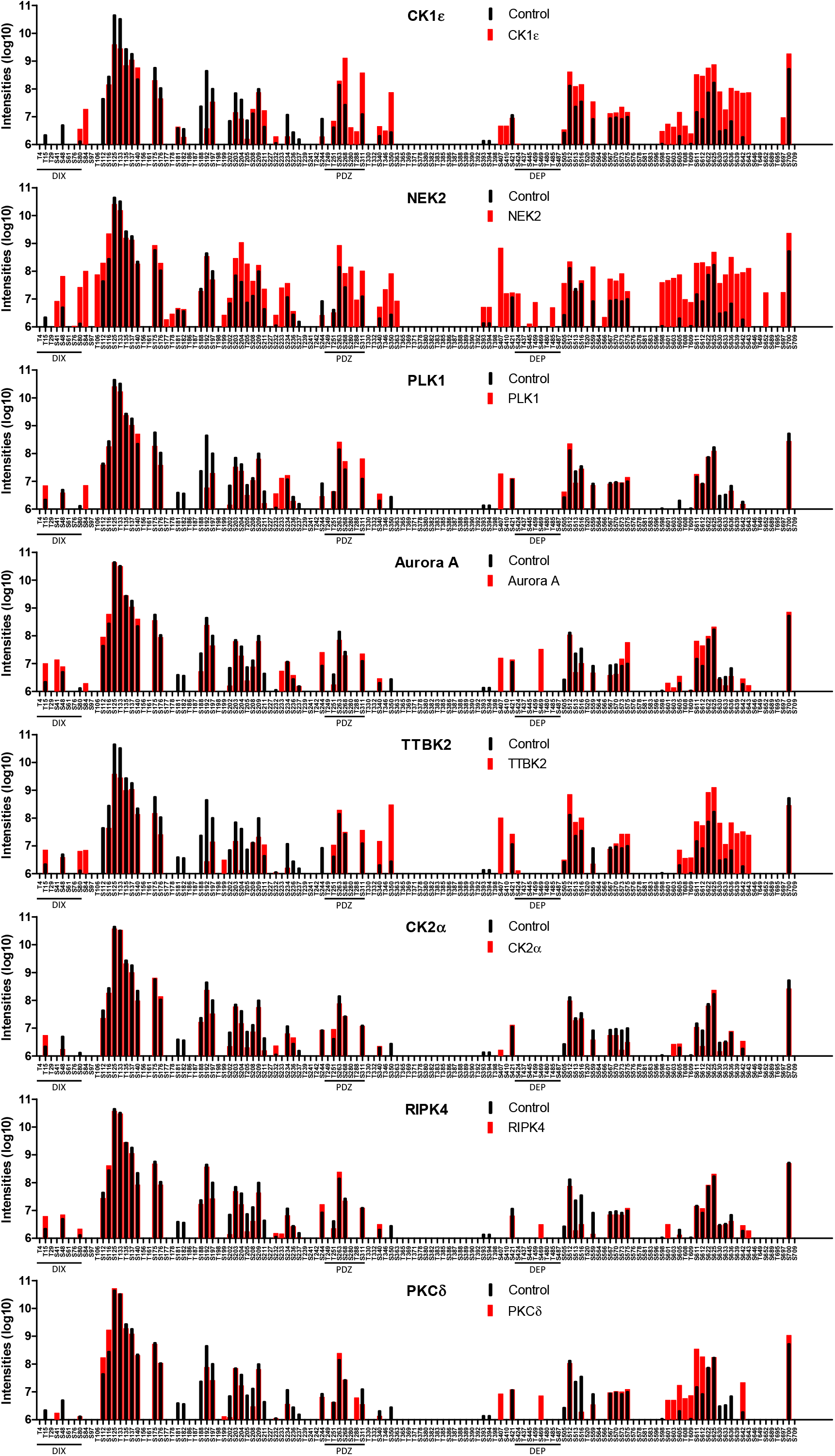
Phosphoplots - phosphorylation barcodes of DVL3 with the individual kinases. Visualization of absolute intensity of phosphorylated peptides corresponding to the individual phosphorylated sites plotted on the primary sequence of DVL3 (only Ser and Thr are shown). Black bars represent a control condition, red bars the intensities in the presence of the kinase. Intensities are plotted on a log10 scale.

In this study we have identified TTBK2 and Aurora A as novel DVL kinases. For TTBK2 we propose a possible negative role in Wnt/β-catenin signaling. TTBK2 is found localized to basal bodies or distal appendages, respectively. Indeed, DVL has been associated with centrosome and basal bodies *(10, 33, 39)*. However, it does not seem to localize to distal appendages of mother centriole, where TTBK2 is recruited by CEP164 to trigger ciliogenesis *(25, 26, 28–31)*. It remains to determined how TTBK2 activity interplays with the phosphorylation of DVL3 by Aurora A, NEK2 and PLK1 and how these kinases together regulate DVL localization and functions in the individual phases of the cell cycle.

## MATERIALS AND METHODS

### Cell Culture, Transfection, and Treatments

HEK293T cells were propagated in DMEM, 10% FCS, 2 mM L-glutamine, 50 units/ml penicillin, 50 units/ml streptomycin. CK1ε inhibitor PF-670462 was used at 10 μM. Cells were seeded on appropriate culture dishes (15 cm diameter dish for IP, 24-well plate for WB, dual luciferase assay, ICC) and the next day were transfected using polyethylenimine (PEI) in a stoichiometry of 3 μl PEI (0.1% w/v in MQ water) per 1 μg of DNA. Cells were harvested for immunoblotting or immunocytofluorescence 24 h after transfection, for immunoprecipitation after 48 h. The following plasmids have been published previously: FLAG-DVL3 *(40)*, CK1ε *(41)*, Myc-NEK2 *(42)*, FLAG-PLK1 *(43)*, Myc-Aurora A *(44)*, HA-CK2α *(45)*, YFP-PKCδ *(46)*, VSV-RIPK4 *(47)* and GFP-TTBK2 WT and KD *(25)*.

### Dual Luciferase TopFlash/Renilla Reporter Assay

For the luciferase reporter assay, cells were transfected with 0.1 μg of Super8X TopFlash construct, 0.1 μg of pRLTKluc (Renilla) luciferase construct and other plasmids as indicated in a 24-well plate and processed 24 h after transfection. For the TopFlash assay, a Promega dual luciferase assay kit was used according to the manufacturer’s instructions. Relative luciferase units of firefly luciferase were measured and normalized to the Renilla luciferase signal.

### Coimmunoprecipitation and Western blotting

For MS/MS-based identification of phosphorylation, HEK293T cells were seeded on 15 cm dishes and transfected with corresponding plasmids (8 μg of DVL3 plasmid plus 8 μg of pCDNA3 or plasmid encoding the kinase per dish) 24 h after seeding. Two ml of ice-cold lysis buffer supplemented with protease inhibitors (Roche Applied Science, 11836145001), phosphatase inhibitors (Calbiochem, 524625), 0.1 mM DTT, and 10 mM N-ethylmaleimide (Sigma E3876) was used for lysis of one 15 cm dish. Lysate was collected after 20 min of lysis on 4 °C and was cleared by centrifugation at 18 000 × g for 20 min. Three μg of anti-FLAG M2 (F1804; Sigma) antibody were used per sample. Samples were incubated with the antibody for 40 min, then 25 μl of G protein-Sepharose beads (GE Healthcare, 17–0618-05) equilibrated in the lysis buffer were added to each sample. Samples were incubated on the carousel overnight and washed 6 times with lysis buffer; finally 40 μl of 2× Laemmli buffer was added, and samples were boiled.

The samples were loaded to 8% SDS-PAGE gels. Electrophoresis was preformed through the stacking gel at 100 V and through the separating gel at 150 with PageRuler Prestained Protein Ladder (Thermofisher, Cat. No. 26620) as a marker. Gels were then process for mass spectrometry or the proteins were transferred to the polyvinylidene difluoride membrane (Merck Millipore, Cat. No. IPVH00010) by Western Blotting (WB). Subsequently, the membrane was blocked in 5% unfatted milk or 3% BSA for 1 hour with shaking. After the blocking step, the membrane was incubated with corresponding primary and secondary antibodies. Proteins were visualized by ECL (Enhanced chemiluminescence; Merck Millipore, Cat. No. WBKLS0500). The signal was detected using Vilber FUSION-SL system. Western blot was quantified using ImageJ software.

The antibodies used were: anti-FLAG M2 (Sigma-Aldrich, #F1804) for WB and IP, anti-CK1□ (Santa Cruz, #sc-6471), anti-GFP (Fitzgerald, #20R-GR-011), HA11 (Covance #MMS-101R), VSV (Sigma-Aldrich #V 5507), c-Myc (Santa Cruz, #sc-40), α-tubulin (Proteintech, 66031-1-Ig). Following phospho-specific antibodies have been published previously – pSer280-DVL3 (pS280) *(10)*, pS643 *(21)*, and pS697 *(10)*. Anti phospho-S192 antibody was prepared by immunizing rabbits by TTSFFDS(p)DEDDST peptide on a service basis by Moravian Biotechnology (http://www.moravian-biotech.com).

### Immunofluorescence

Cells were seeded onto glass coverslips and next day transfected according to the scheme. 24 hours post transfection medium was removed, cells were washed by PBS and fixed by 4% PFA for 10 minutes or ice-cold MetOH (5 min/-20 °C, Fig. 1B). Coverslips were then washed in PBS and incubated with primary antibodies for 1 h, washed three times with PBS and then incubated with secondary antibodies conjugated to Alexa Fluor 488 (Invitrogen A11001) or/and Alexa Fluor 594 (Invitrogen A11058), washed with PBS and stained with DAPI (1:5000); all coverslips were mounted on microscopic slides. Cells were then visualized on Olympus IX51 fluorescent microscope using 40× air or 100× oil objectives and/or Olympus Fluoview 500 confocal laser scanning microscope IX71 using 100× oil objective. 200 positive cells per experiment (N=3) were analyzed and scored according to their phenotype into two categories (punctae/even). The antibodies used were as follows: anti-FLAG M2 (Sigma-Aldrich, #F1804), anti-DVL3 (Santa Cruz, #sc-8027), anti-CK1□ (Santa Cruz, #sc-6471), HA11 (Covance #MMS-101R), VSV (Sigma-Aldrich #V 5507), c-Myc (Santa Cruz, #sc-40) and anti-GFP (Fitzgerald, #20R-GR-011), CAP350 *(48)*, TTBK2 (Sigma-Aldrich, #HPA018113). Images presented in Figure 1B were acquired using DeltaVision-Elite system (Applied Precision/GE) with a 100×/1.4 Apo plan oil immersion objective. Image stacks were taken with a z distance of 0.2 μm, deconvolved (conservative ratio, three cycles), and projected as maximal intensity images by using SoftWoRX (Applied Precision/GE).

### Mass spectrometry

#### In gel digestion

Immunoprecipitates were separated on SDS-PAGE gel electrophoresis, fixed with acetic acid in methanol and stained with Coomassie brilliant blue for 1 hour. Corresponding 1D bands were excised. After destaining, the proteins in gel pieces were incubated with 10 mM DTT at 56 °C for 45 min. After removal of DTT excess samples were incubated with 55 mM IAA at room temperature in darkness for 30 min, then alkylation solution was removed and gel pieces were hydrated for 45 min at 4 °C in digestion solution (5 ng/μl trypsin, sequencing grade, Promega, in 25 mM AB). The trypsin digestion proceeded for 2 hours at 37 °C on Thermomixer (750 rpm; Eppendorf). Subsequently, the tryptic digests were cleaved by chymotrypsin (5 ng/μl, sequencing grade, Roche, in 25 mM AB) for 2 hours at 37 °C. Digested peptides were extracted from gels using 50% ACN solution with 2.5% formic acid (FA) and concentrated in speedVac concentrator (Eppendorf). The aliquot (1/10) of concentrated sample was transferred to LC-MS vial with already added polyethylene glycol (PEG; final concentration 0.001%, *(49)* and directly analyzed by LC-MS/MS for protein identification.

#### Phosphopeptide enrichment

The rest of the sample (9/10) was used for phosphopeptide analysis. Sample was diluted with acidified acetonitrile solution (80% ACN, 2% FA). Phosphopeptides were enriched using Pierce Magnetic Titanium Dioxide Phosphopeptide Enrichment Kit (Thermo Scientific, Waltham, Massachusetts, USA) according to manufacturer protocol and eluted into LC-MS vial with already added PEG (final concentration 0.001%). Eluates were concentrated under vacuum and then dissolved in water and 0.6 μl of 5% FA to get 12 μl of peptide solution before LC-MS/MS analysis.

#### LC-MS/MS analysis

LC-MS/MS analyses of peptide mixture were done using RSLCnano system connected to Orbitrap Elite hybrid spectrometer (Thermo Fisher Scientific) with ABIRD (Active Background Ion Reduction Device; ESI Source Solutions) and Digital PicoView 550 (New Objective) ion source (tip rinsing by 50% acetonitrile with 0.1% formic acid) installed. Prior to LC separation, peptide samples were online concentrated and desalted using trapping column (100 μm × 30 mm) filled with 3.5 μm X-Bridge BEH 130 C18 sorbent (Waters). After washing of trapping column with 0.1% FA, the peptides were eluted (flow 300 nl/min) from the trapping column onto Acclaim Pepmap100 C18 column (3 μm particles, 75 μm × 500 mm; Thermo Fisher Scientific) by 65 min long gradient. Mobile phase A (0.1% FA in water) and mobile phase B (0.1% FA in 80% acetonitrile) were used in both cases. The gradient elution started at 1% of mobile phase B and increased from 1% to 56% during the first 50 min (30% in the 35^th^ and 56% in 50^th^ min), then increased linearly to 80% of mobile phase B in the next 5 min and remained at this state for the next 10 min. Equilibration of the trapping column and the column was done prior to sample injection to sample loop. The analytical column outlet was directly connected to the Digital PicoView 550 ion source.

MS data were acquired in a data-dependent strategy selecting up to top 10 precursors based on precursor abundance in the survey scan (350–2000 m/z). The resolution of the survey scan was 60 000 (400 m/z) with a target value of 1×10^6^ ions, one microscan and maximum injection time of 200 ms. High resolution (15 000 at 400 m/z) HCD MS/MS spectra were acquired with a target value of 50 000. Normalized collision energy was 32% for HCD spectra. The maximum injection time for MS/MS was 500 ms. Dynamic exclusion was enabled for 45 s after one MS/MS spectra acquisition and early expiration was disabled. The isolation window for MS/MS fragmentation was set to 2 m/z.

#### Data analysis

The analysis of the mass spectrometric RAW data was carried out using the Proteome Discoverer software (Thermo Fisher Scientific; version 1.4) with in-house Mascot (Matrixscience; version 2.4.1) search engine utilization. MS/MS ion searches were done against in-house database containing expected protein of interest with additional sequences from cRAP database (downloaded from http://www.thegpm.org/crap/). Mass tolerance for peptides and MS/MS fragments were 7 ppm and 0.03 Da, respectively. Oxidation of methionine, deamidation (N, Q) and phosphorylation (S, T, Y) as optional modification, carbamidomethylation of C as fixed modification, TrypChymo enzyme specifity and three enzyme miss cleavages were set for all searches. The phosphoRS (version 3.1) feature was used for preliminary phosphorylation localization. Final localization of all phosphorylations (including those with ambiguous localization) was performed by manual evaluation of the fragmentation spectra of the individual phosphopeptides. Based on the presence of individual fragments in the peptide sequence, it was decided whether the localization was accurate or not.

Quantitative information was assessed and manually validated in Skyline software (Skyline daily 3.6.1.10230). Normalization of the data was performed using the set of phosphopeptide standards (added to the sample prior phosphoenrichment step; MS PhosphoMix 1, 2, 3 Light, Sigma) and by non-phosphorylated peptides identified in direct analyses.

All quantitative data (peptide intensities) were processed by two approaches (see Fig. 3A). In the first approach (pipeline #1 and #3) that often resulted in the formation of phosphorylated clusters all identified phosphorylated peptides were considered. This has resulted in three categories of identification: (i) one peptide or set of peptides with one clearly localized phosphorylated site, (ii) one or set of overlapping peptides covering sequence region with two or more clearly localized phosphorylated sites (phosphorylated sites separated by a comma in the Figures) and (iii) one or set of overlapping peptides covering sequence region with two or more phosphorylated sites, but some of them are not clearly localized (phosphorylated sites separated by a dash in the Figures). Sum of intensities of phosphorylated peptides (includes different peptide sequences, charges and peptides with other modifications) was calculated for each cluster. In case of direct analysis (pipeline #1), we calculated site occupancies as percentage ratio of phosphorylated peptide intensity (summed if more than one) to total intensity (summed intensities of phosphorylated peptide(s) + corresponding non-phosphorylated peptide(s)). Sites/clusters with the site occupancy >5 % at least in one experiment are shown in Fig. 4. For pipeline #3, the difference between the phosphorylated peptide summed intensity in the kinase-induced sample and the control (in log10 scale) was calculated for each replicate and the mean from all three replicates was used for the heat map (see in Supplementary Fig. 2). In the second approach (pipeline #2), only phosphorylated peptides with the clearly localized phosphorylation site (based on manual inspection) were considered. In case of the multiphosphorylated peptides the total intensity of the peptide was assigned to each phosphorylated site. Sum of intensities of phosphorylated peptides was calculated for each phosphorylated site. The heat map (Fig. 5) was build-up in the same way as for pipeline #3 described above. For the production of phosphoplots the sum of intensities for each phosphorylated site (pipeline #2) were log-transformed (log10) and average from 3 replicates was calculated (values under the detection limit were not included in the calculation).

### Numerical data & Statistics

#### Ouster analysis

Data of absolute peptide intensities were log-transformed (log10) because of their log-normal distribution. Log-transformed data of each replicate were standardized to control (kinase subtracted from “no kinase” control) and averaged from 3 replicates. Cluster analysis was applied both for kinases and for phosphosites; Ward’s minimum variance method based on the Lance-Williams recurrence and Euclidean distance were used. Circular plot accompanied by heat map was used for visualization of complex relation among phosphosites and kinases. All analyses were performed using R software.

#### Other analyses

One-way ANOVA and Tukey Post tests were calculated by GraphPad Prism (GraphPad Software Inc.).

## SUPPLEMENTARY MATERIALS

Fig. S1. SDS-PAGE gels from three independent experiments.

Fig. S2. Phosphorylated clusters map of DVL3.

Fig. S3. Cluster analysis of the individual phosphorylated sites/clusters and individual kinases.

Table S1. Intensities of the individual phosphorylated sites (raw data).

Table S2. Sequence coverage of DVL3 in the individual experiments.

## Acknowledgements

We would like to thank Erich Nigg (Biozentrum, Basel) for sharing reagents, Lumir Krejci for access to DV Elite and Lucie Smyčková, Lenka Bryjová and Nad’a Bílá for excellent assistance. Funding: The work was supported by grants from the Czech Science Foundation (GA17-16680S, GA18-17658S, 19-28347X) to VB and grants from Czech Science Foundation (16-03269Y, 19-05244S); Swiss National Science Foundation (IZ11Z0_166533); and Follow up research fund from Federation of Biochemical and Biophysical Societies (FEBS) to LC. OB was supported by funds from the Faculty of Medicine MU to junior researcher. MK was supported by Brno Ph.D. Talent Scholarship – Funded by the Brno City Municipality. CIISB research infrastructure project LM2015043 funded by MEYS CR is gratefully acknowledged for the financial support of the LC-MS/MS measurements at the Proteomics Core Facility.

## Author contributions

OB, MK, MM, LC and MR performed all cell-based assays, KH prepared samples for MS analysis and processed the data, DP performed LC-MS/MS analyses, PO performed the statistical analyses, VB, KH, DP, LC and ZZ planned the experiments, interpreted the data and wrote the manuscript. All authors have approved the final version.

## Data and materials availability

Proteomic data are available via PRIDE.

