## Supplemental File for "Comparative phosphorylation map of Dishevelled3 (DVL3)"

**Supplementary Figure 1. SDS-PAGE gels from three independent experiments.**

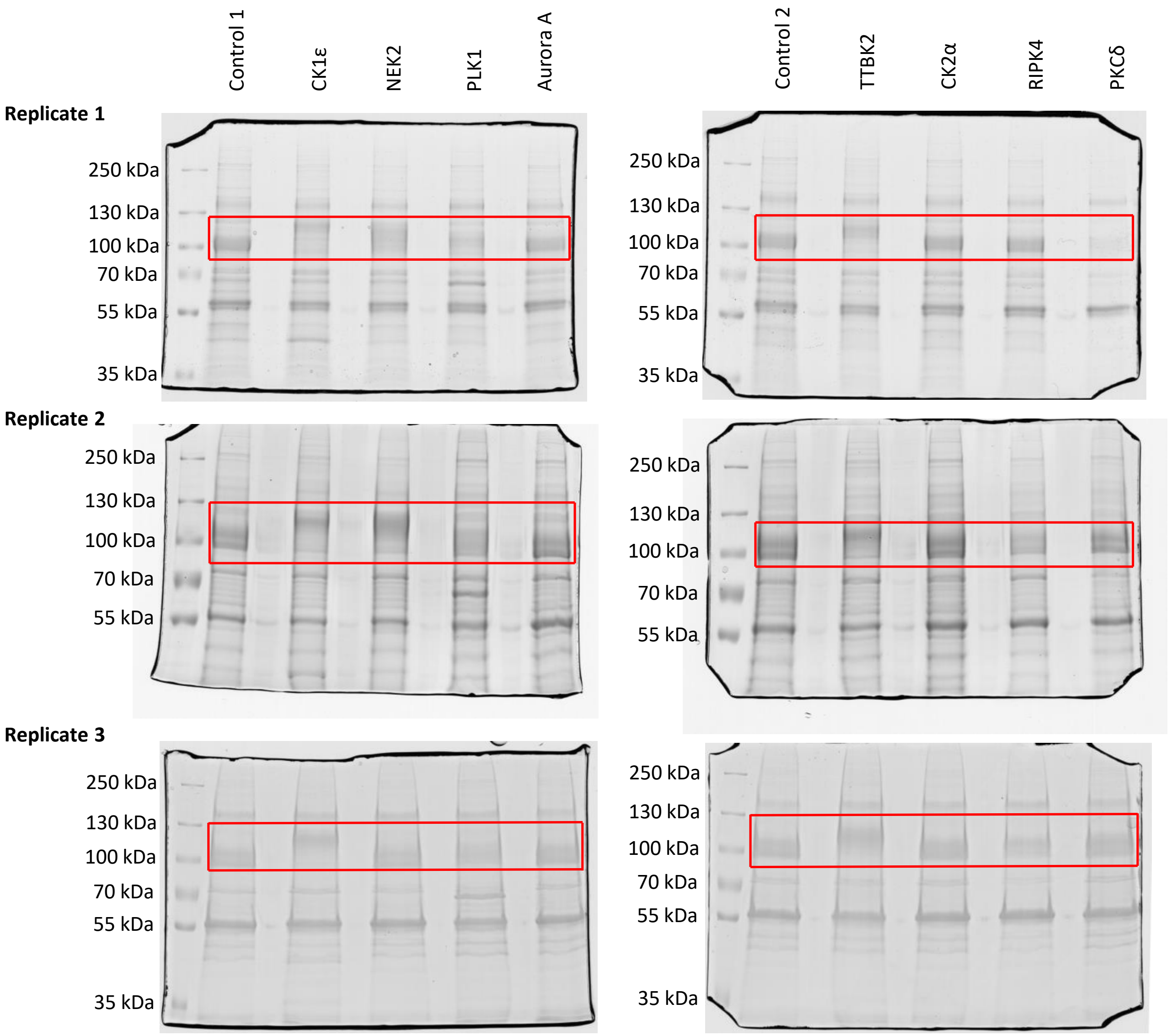


FLAG-DVL3 was overexpressed in HEK293, with or without the studied kinase, immunoprecipitated using the anti-FLAG antibody, separated on SDS-PAGE and stained with Coomassie Brilliant Blue. In red boxes are parts that were cut out.

**Supplementary Figure 2. Map of the phosphorylated clusters of DVL3.**

| DVL3 | Mean intensity of control | Relative change of phosphopeptide intensities | | | | | | | | |
|---|---|---|---|---|---|---|---|---|---|---|
| | | Control | CK1ε | NEK2 | PLK1 | Aurora A | TTBK2 | CK2α | RIPK4 | PKCδ |
| S76, S80, S84 | 6.17 | 2 | 3 | 3 | 3 | 2 | 3 | 2 | 2 | 1 |
| T106–S140 | 10.83 | 3 | 3 | 3 | 3 | 3 | 3 | 3 | 3 | 3 |
| S175, S176, S177 | 8.77 | 3 | 3 | 3 | 3 | 3 | 3 | 3 | 3 | 3 |
| T178–S182 | 6.57 | 2 | 1 | 2 | 1 | 1 | 1 | 1 | 0 | 0 |
| S188, S192, S197 | 8.16 | 3 | 3 | 3 | 3 | 3 | 3 | 3 | 3 | 3 |
| S202–S209 | 8.28 | 3 | 3 | 3 | 3 | 3 | 3 | 3 | 3 | 3 |
| S232–S237 | 7.26 | 3 | 2 | 3 | 3 | 3 | 3 | 3 | 3 | 3 |
| T251, Y257 | 6.36 | 1 | 1 | 1 | 1 | 1 | 1 | 1 | 1 | 1 |
| S393, S394 | 6.24 | 1 | 0 | 1 | 0 | 0 | 0 | 0 | 0 | 0 |
| S407, S410 | 6.06 | 1 | 1 | 1 | 1 | 1 | 1 | 1 | 1 | 1 |
| S505–S516 | 7.84 | 3 | 3 | 3 | 2 | 3 | 3 | 2 | 2 | 2 |
| S559–S575 | 7.63 | 3 | 3 | 3 | 3 | 3 | 3 | 3 | 3 | 3 |
| S598–S612 | 7.13 | 3 | 3 | 3 | 2 | 3 | 3 | 3 | 2 | 3 |
| S622, S625, S630 | 8.17 | 3 | 3 | 3 | 3 | 3 | 3 | 3 | 3 | 3 |
| S633–S643 | 6.88 | 3 | 3 | 3 | 3 | 3 | 3 | 3 | 3 | 2 |

ND (6.00) 11.00 -1.50 2.00

All identified phosphorylated clusters obtained from the pipeline #3 are visualized as a heatmap. Mean absolute intensities of phosphorylation clusters in the control (DVL3 without exogenous kinase; six replicates) are expressed in the shades of blue. Numbers indicate decadic logarithm of the mean of 6 control samples.

Nine columns represent heat map of relative change of phosphorylated peptide intensities (in log10 scale) obtained for individual kinases (relative to control). Numbers in the heat map fields (0, 1, 2, 3) indicate the number of experimental replicates with the positive identification of the given phosphorylated site.

**Supplementary Figure 3. Cluster analysis of the individual phosphorylated sites and individual kinases.**

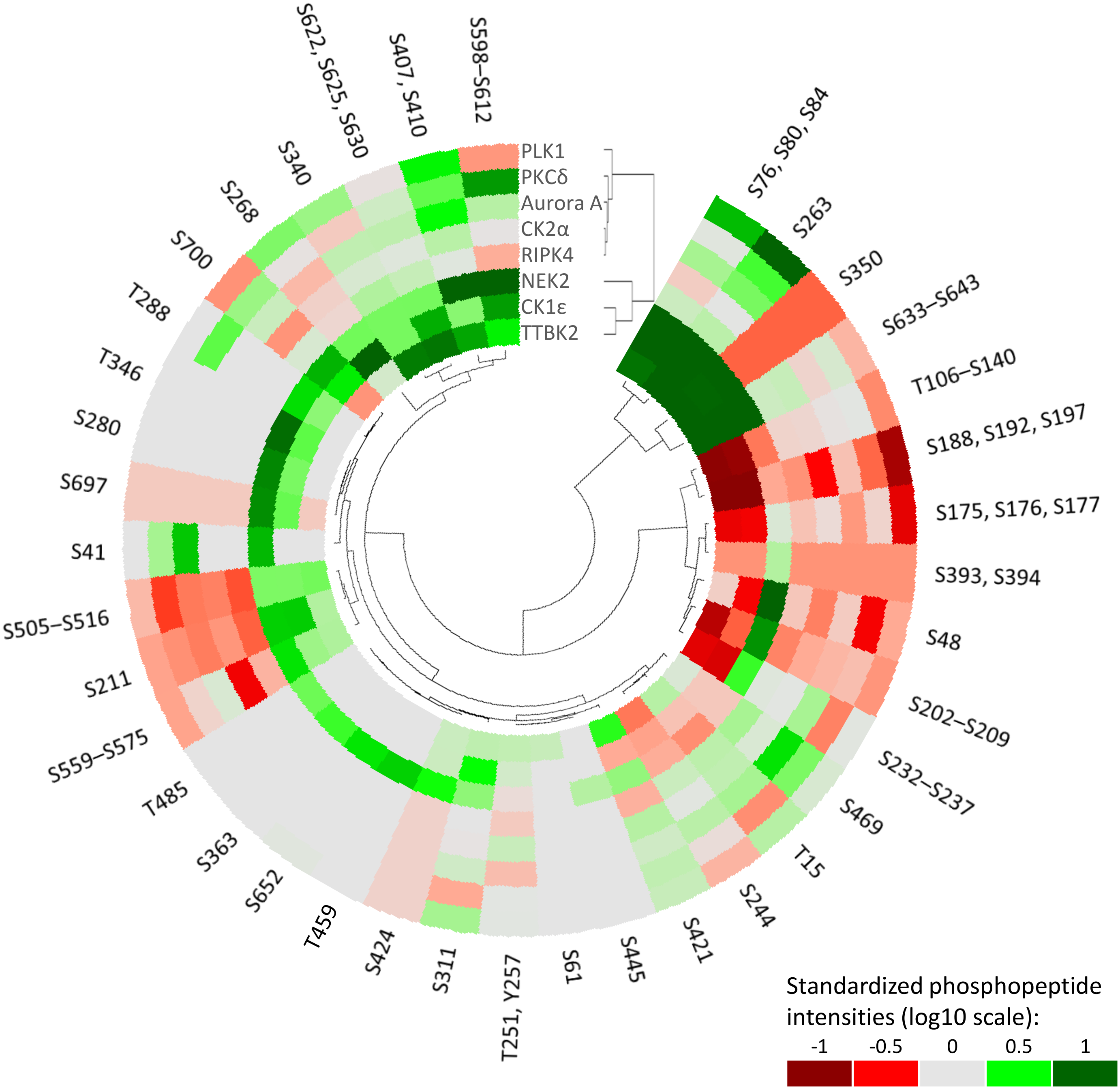


Mean phosphorylated peptide intensities (in log10 scale) standardized to control are used in the circular plot. Peptides expressed more intensively compared to control are in the shades of green, whereas less expressed peptides are in red. Dendrograms both for the kinases and the phosphosites are shown.

### Supplementary Table1. Phosphosite intensities in whole experiment

| Phosphorylated sites | Control 1 | | CK1ε | | NEK2 | | PLK1 | | Aurora A | | Control 2 | | TTBK2 | | CK2α | | RIPK4 | | PKCδ | |
|---|---|---|---|---|---|---|---|---|---|---|---|---|---|---|---|---|---|---|---|---|
| | Direct | Enriched | Direct | Enriched | Direct | Enriched | Direct | Enriched | Direct | Enriched | Direct | Enriched | Direct | Enriched | Direct | Enriched | Direct | Enriched | Direct | Enriched |
| T15 | 7.85 | 6.20 | 8.00 | 6.15 | 7.96 | ND | 8.15 | 6.44 | 7.89 | 6.49 | 7.95 | 6.32 | 7.87 | 6.44 | 7.93 | 6.41 | 7.72 | 6.42 | 7.95 | ND |
| S41 | ND | ND | ND | ND | 7.38 | 6.72 | ND | ND | 7.16 | 6.67 | ND | ND | ND | ND | ND | ND | 6.39 | ND | ND | 6.24 |
| S48 | ND | 6.65 | ND | ND | ND | 7.61 | ND | 6.36 | ND | 6.46 | ND | 6.42 | ND | 6.47 | ND | 6.24 | ND | 6.44 | ND | ND |
| S61 | ND | ND | ND | ND | ND | ND | ND | ND | ND | ND | ND | ND | ND | 6.12 | ND | ND | ND | ND | ND | ND |
| S76 | ND | ND | ND | ND | ND | 6.17 | ND | ND | ND | ND | ND | ND | ND | ND | ND | ND | ND | ND | ND | ND |
| S80 | ND | 6.23 | 6.23 | 6.46 | 6.92 | 6.97 | 6.36 | 6.09 | 6.44 | 6.13 | 6.35 | 6.12 | 6.40 | 6.65 | ND | 6.10 | ND | 6.28 | ND | 6.19 |
| S84 | 6.42 | ND | 7.33 | 7.21 | 8.16 | 7.75 | 7.12 | 6.84 | 6.35 | 6.28 | ND | ND | 6.36 | 6.77 | 6.10 | ND | ND | ND | ND | ND |
| T106 | ND | ND | ND | ND | ND | 7.67 | ND | ND | ND | ND | ND | ND | ND | ND | ND | ND | ND | ND | ND | ND |
| S112 | ND | 6.71 | ND | ND | ND | 7.62 | ND | 6.69 | ND | 6.81 | ND | 6.69 | ND | ND | ND | 6.61 | ND | 6.64 | ND | 6.90 |
| S116 | ND | 8.42 | ND | 8.15 | ND | 9.20 | ND | 8.15 | ND | 8.77 | ND | 8.31 | ND | 7.59 | ND | 8.21 | ND | 8.55 | ND | 8.95 |
| S125 | 9.68 | 10.64 | 8.95 | 9.56 | 9.77 | 10.33 | 9.81 | 10.33 | 9.55 | 10.60 | 9.77 | 10.57 | 8.70 | 9.57 | 9.54 | 10.56 | 9.73 | 10.56 | 9.66 | 10.62 |
| T133 | 9.53 | 10.35 | 8.83 | 9.27 | 9.45 | 9.96 | 9.75 | 10.11 | 9.42 | 10.32 | 9.63 | 10.28 | 8.60 | 9.31 | 9.43 | 10.32 | 9.60 | 10.29 | 9.55 | 10.34 |
| T135 | 9.29 | 9.43 | 8.75 | 7.95 | 9.31 | 9.17 | 9.58 | 9.36 | 9.17 | 9.43 | 9.39 | 9.36 | 8.40 | 7.97 | 9.21 | 9.12 | 9.38 | 9.42 | 9.35 | 9.28 |
| S137 | 6.36 | 9.23 | 6.20 | 8.97 | 6.69 | 9.07 | 6.30 | 9.00 | ND | 9.03 | 6.43 | 9.18 | ND | 8.80 | 6.38 | 8.96 | 6.09 | 9.03 | ND | 9.07 |
| S140 | ND | 7.75 | ND | 7.78 | ND | 7.62 | ND | 7.88 | ND | 7.78 | ND | 7.55 | ND | 7.51 | ND | 7.43 | ND | 7.34 | ND | 7.51 |
| S175 | ND | 8.80 | ND | 8.17 | ND | 8.76 | ND | 8.11 | ND | 8.47 | ND | 8.62 | ND | 8.13 | ND | 8.68 | ND | 8.61 | ND | 8.66 |
| S176 | 8.14 | 7.46 | 7.20 | 7.18 | 8.19 | 7.58 | 7.65 | 7.16 | 7.94 | 7.41 | 7.87 | 7.46 | 6.88 | 7.04 | 7.91 | 7.53 | 7.58 | 7.40 | 8.01 | 7.47 |
| S177 | ND | 6.10 | ND | ND | ND | 6.25 | ND | ND | ND | ND | ND | ND | ND | ND | ND | ND | ND | ND | ND | ND |
| T178 | ND | ND | ND | ND | ND | 6.31 | ND | ND | ND | ND | ND | ND | ND | ND | ND | ND | ND | ND | ND | ND |
| S181 | ND | 6.31 | ND | 6.37 | ND | 6.45 | ND | ND | ND | ND | ND | 6.39 | ND | ND | ND | 6.08 | ND | ND | ND | ND |
| S182 | ND | 6.31 | ND | 6.25 | ND | 6.54 | ND | 6.09 | ND | 6.10 | ND | 6.53 | ND | 6.10 | ND | 6.08 | ND | ND | ND | ND |
| S188 | ND | 6.64 | ND | 6.16 | ND | 6.59 | ND | ND | ND | 6.40 | ND | 6.58 | ND | ND | ND | 6.56 | ND | 6.57 | ND | ND |
| S192 | 9.33 | 7.05 | 6.69 | 6.35 | 9.06 | 7.00 | 7.55 | 6.41 | 9.12 | 6.95 | 9.39 | 7.03 | 6.89 | 6.30 | 9.10 | 6.95 | 9.17 | 7.01 | 9.17 | 6.79 |
| S197 | 7.16 | 7.95 | 6.18 | 7.08 | 6.93 | 7.62 | ND | 7.21 | 7.15 | 7.59 | 7.38 | 7.75 | ND | 6.98 | 6.55 | 7.30 | ND | 7.38 | 6.72 | 7.32 |
| S199 | ND | ND | ND | ND | ND | 6.30 | ND | ND | ND | ND | ND | ND | ND | 6.32 | ND | ND | ND | ND | ND | 6.20 |
| S202 | ND | 6.65 | ND | ND | ND | 6.66 | ND | 6.18 | ND | 6.22 | ND | 6.58 | ND | 6.08 | ND | 6.34 | ND | 6.20 | ND | 6.17 |
| S203 | 8.05 | 7.47 | 6.53 | 6.77 | 8.78 | 8.11 | 7.88 | 7.30 | 7.86 | 7.35 | 8.17 | 7.43 | 6.33 | 6.76 | 8.15 | 7.29 | 8.06 | 7.21 | 7.45 | 7.35 |
| S204 | 7.46 | 7.58 | 6.44 | 6.75 | 9.16 | 9.01 | 7.01 | 7.34 | 6.81 | 7.26 | 7.04 | 7.50 | ND | 6.15 | 7.16 | 7.15 | 6.85 | 7.08 | 6.80 | 7.17 |
| T205 | ND | 6.46 | ND | 6.23 | 7.50 | 7.27 | ND | 6.33 | ND | 6.29 | ND | 6.44 | ND | ND | ND | 6.26 | ND | 6.24 | ND | ND |
| S208 | ND | 7.08 | ND | 7.18 | ND | 7.55 | ND | 6.97 | ND | 6.86 | ND | 7.00 | ND | 6.91 | ND | 6.70 | ND | 6.58 | ND | 6.43 |
| S209 | 7.97 | 7.72 | 7.30 | 7.72 | 8.03 | 7.87 | 7.83 | 7.67 | 7.79 | 7.59 | 7.76 | 7.73 | 6.77 | 7.30 | 7.63 | 7.61 | 7.40 | 7.32 | 7.07 | 7.50 |
| S211 | ND | 6.38 | ND | 7.04 | ND | 7.02 | ND | 6.23 | ND | 6.10 | ND | 6.36 | ND | 6.65 | ND | 6.22 | ND | ND | ND | 6.12 |
| S232 | ND | 6.19 | ND | 6.25 | ND | 6.30 | ND | 6.35 | ND | 6.10 | ND | 6.16 | ND | 6.17 | ND | 6.28 | ND | 6.21 | ND | 6.17 |
| S233 | ND | 6.11 | ND | ND | ND | 7.32 | ND | 6.85 | ND | 6.60 | ND | 6.13 | ND | 6.16 | ND | 6.13 | ND | 6.21 | ND | ND |
| S234 | ND | 6.56 | ND | 6.26 | ND | 7.46 | ND | 7.04 | ND | 6.75 | ND | 6.44 | ND | 6.23 | ND | 6.43 | ND | 6.43 | ND | 6.35 |
| S236 | ND | 6.27 | ND | ND | ND | 6.35 | ND | 6.26 | ND | 6.50 | ND | 6.42 | ND | ND | ND | 6.48 | ND | 6.39 | ND | 6.10 |
| S237 | ND | 6.27 | ND | ND | ND | ND | ND | ND | ND | 6.22 | ND | 6.13 | ND | ND | ND | 6.13 | ND | 6.17 | ND | ND |
| S244 | ND | 6.52 | ND | 6.25 | ND | 6.30 | ND | 6.31 | ND | 6.63 | ND | 6.39 | ND | 6.14 | ND | 6.47 | ND | 6.56 | ND | 6.43 |
| T251 | ND | 6.32 | ND | 6.44 | ND | 6.33 | ND | 6.37 | ND | 6.24 | ND | 6.40 | ND | 6.50 | ND | 6.48 | ND | 6.28 | ND | 6.37 |
| Y257 | ND | 6.32 | ND | 6.44 | ND | 6.33 | ND | 6.37 | ND | 6.24 | ND | 6.40 | ND | 6.50 | ND | 6.48 | ND | 6.28 | ND | 6.37 |
| S263 | ND | 6.92 | 6.88 | 8.10 | 9.03 | 8.82 | 8.54 | 8.38 | 6.50 | 7.30 | 6.20 | 6.94 | 6.94 | 7.96 | ND | 6.93 | ND | 7.10 | ND | 7.39 |
| S268 | 6.95 | 7.35 | 8.93 | 9.05 | 7.06 | 7.70 | 7.23 | 7.67 | 6.68 | 7.19 | 6.68 | 7.30 | 6.56 | 7.40 | 6.19 | 7.26 | 6.68 | 7.28 | 6.57 | 7.31 |
| S280 | ND | ND | 7.03 | 6.36 | 7.70 | 6.88 | 6.34 | ND | ND | ND | ND | ND | ND | ND | ND | ND | ND | ND | ND | ND |
| T288 | ND | ND | ND | 6.31 | ND | 6.59 | ND | ND | ND | ND | ND | ND | ND | ND | ND | ND | ND | ND | ND | 6.42 |
| S311 | 6.38 | 6.52 | 7.81 | 7.02 | 6.71 | 6.83 | 6.65 | 6.76 | 6.39 | 6.61 | ND | 6.52 | 6.81 | 6.68 | 6.35 | 6.51 | ND | 6.52 | ND | 6.34 |
| S340 | 6.07 | 6.34 | 6.81 | 6.63 | 7.13 | 6.63 | 7.08 | 6.54 | 6.49 | 6.41 | 6.36 | 6.19 | 7.56 | 7.17 | 6.35 | 6.32 | 7.01 | 6.47 | 6.27 | 6.18 |
| T346 | ND | ND | ND | 6.42 | ND | 6.96 | ND | ND | ND | ND | ND | ND | ND | ND | ND | ND | ND | ND | ND | ND |
| S350 | 6.70 | 6.27 | 7.78 | 7.36 | 7.79 | 7.37 | ND | ND | 6.59 | ND | 6.67 | 6.46 | 8.04 | 8.42 | 6.77 | ND | ND | ND | ND | ND |
| S363 | ND | ND | ND | ND | ND | 6.47 | ND | ND | ND | ND | ND | ND | ND | ND | ND | ND | ND | ND | ND | ND |
| S393 | ND | 6.24 | ND | ND | ND | 6.40 | ND | ND | ND | ND | ND | 6.15 | ND | ND | ND | ND | ND | ND | ND | ND |
| S394 | ND | 6.24 | ND | ND | ND | 6.40 | ND | ND | ND | ND | ND | 6.15 | ND | ND | ND | ND | ND | ND | ND | ND |
| S407 | 6.46 | ND | ND | 6.38 | 8.73 | 7.10 | 7.80 | 6.58 | 6.68 | 6.56 | 6.42 | 6.11 | 8.31 | 6.83 | ND | 6.23 | ND | 6.08 | 6.42 | 6.47 |
| S410 | ND | ND | ND | 6.38 | ND | 6.56 | ND | ND | ND | ND | ND | ND | ND | ND | ND | ND | ND | ND | ND | ND |
| S421 | 6.36 | 7.02 | ND | 6.75 | 6.37 | 7.21 | 6.54 | 7.04 | 6.38 | 7.00 | 6.34 | 6.80 | 6.54 | 7.39 | 6.23 | 7.10 | 6.42 | 6.75 | ND | 7.06 |
| S424 | ND | ND | ND | 6.17 | 6.44 | 6.55 | ND | ND | 6.27 | ND | 6.20 | 6.11 | 6.21 | 6.20 | ND | ND | ND | ND | ND | ND |
| S445 | ND | ND | ND | ND | 6.42 | 6.19 | ND | ND | ND | ND | ND | ND | ND | ND | ND | ND | ND | ND | ND | ND |
| T459 | ND | ND | ND | 6.01 | ND | 6.64 | ND | ND | ND | ND | ND | ND | ND | ND | ND | ND | ND | ND | ND | ND |
| S469 | 6.50 | 6.07 | ND | ND | 6.93 | ND | 6.43 | 6.14 | 6.77 | 6.67 | 6.40 | 6.11 | 6.41 | 6.13 | 6.45 | 6.14 | 6.56 | 6.32 | 6.47 | 6.45 |
| T485 | ND | ND | ND | ND | 6.61 | 6.39 | ND | ND | ND | ND | ND | ND | ND | ND | ND | ND | ND | ND | ND | ND |
| S505 | ND | 6.50 | 6.90 | 6.47 | 7.57 | 7.32 | 7.12 | 6.44 | ND | 6.03 | 6.32 | 6.03 | ND | 6.44 | ND | ND | ND | ND | ND | ND |
| S512 | ND | 7.47 | 7.11 | 7.94 | ND | 7.85 | ND | 7.47 | ND | 7.21 | ND | 7.41 | 6.90 | 7.90 | ND | 7.35 | ND | 7.03 | ND | 7.33 |
| S513 | ND | 6.61 | 7.11 | 6.86 | ND | 6.59 | ND | 6.47 | ND | 6.39 | ND | 6.61 | 6.90 | 6.78 | ND | 6.59 | ND | 6.25 | ND | ND |
| S516 | 7.34 | 6.66 | 7.80 | 7.35 | 6.68 | 7.19 | ND | 7.08 | ND | 6.79 | 7.21 | 7.06 | 6.93 | 7.36 | 6.55 | 6.61 | ND | 6.33 | 6.67 | 6.25 |
| S559 | ND | 6.67 | ND | 7.10 | 6.76 | 7.28 | ND | 6.68 | ND | 6.55 | ND | 6.36 | ND | 6.27 | ND | 6.35 | ND | 6.21 | ND | 6.34 |
| S566 | ND | ND | ND | ND | ND | 6.27 | ND | ND | ND | ND | ND | ND | ND | ND | ND | ND | ND | ND | ND | ND |
| S567 | ND | 6.67 | ND | 6.81 | ND | 7.17 | ND | 6.69 | ND | 6.48 | ND | 6.68 | ND | 6.70 | ND | 6.41 | ND | 6.63 | ND | 6.71 |
| S570 | ND | 6.69 | ND | 6.82 | ND | 7.14 | ND | 6.72 | ND | 6.49 | ND | 6.70 | ND | 6.79 | ND | 6.41 | ND | 6.64 | ND | 6.73 |
| S573 | ND | 6.79 | ND | 7.02 | ND | 7.28 | ND | 6.67 | ND | 6.63 | ND | 6.60 | ND | 6.99 | ND | 6.26 | ND | 6.52 | ND | 6.70 |
| S575 | ND | 6.79 | ND | 6.61 | ND | 6.58 | ND | 6.58 | ND | 7.29 | ND | 6.59 | ND | 6.63 | ND | 6.43 | ND | 6.72 | ND | 6.52 |
| S598 | ND | 6.14 | ND | 6.32 | ND | 6.69 | ND | ND | ND | ND | ND | 6.19 | ND | ND | ND | ND | ND | 6.14 | ND | ND |
| S601 | ND | ND | ND | 6.61 | ND | 7.15 | ND | ND | ND | 6.32 | ND | 6.04 | ND | ND | ND | ND | ND | 6.32 | ND | 6.39 |
| S603 | ND | ND | ND | 6.51 | ND | 7.19 | ND | ND | ND | 6.17 | ND | 6.19 | ND | ND | ND | 6.30 | ND | 6.14 | ND | 6.39 |
| S605 | ND | 6.09 | ND | 6.73 | ND | 7.73 | ND | 6.05 | ND | 6.33 | ND | 6.39 | ND | 6.77 | ND | 6.30 | ND | 6.18 | ND | 6.68 |
| T608 | ND | ND | ND | 6.38 | ND | 6.75 | ND | ND | ND | 6.11 | ND | ND | ND | 6.44 | ND | ND | ND | ND | ND | 6.50 |
| T609 | ND | 6.27 | ND | 6.29 | ND | 6.45 | ND | ND | ND | 6.17 | ND | ND | ND | 6.35 | ND | 6.13 | ND | ND | ND | 6.45 |
| S611 | 6.07 | 7.11 | 6.54 | 7.60 | 7.37 | 7.09 | 6.44 | 6.89 | 6.60 | 7.10 | ND | 6.45 | ND | 7.25 | ND | 6.78 | 6.28 | 6.78 | 6.87 | 7.47 |
| S612 | ND | 6.61 | ND | 7.29 | ND | 7.59 | ND | 6.46 | ND | 6.71 | ND | 6.59 | ND | 6.73 | ND | 6.27 | ND | 6.65 | ND | 6.91 |
| S622 | 7.01 | 7.88 | 8.66 | 8.73 | 7.77 | 8.08 | 7.36 | 7.77 | 7.43 | 7.90 | 7.30 | 7.75 | 8.75 | 8.88 | 7.02 | 7.65 | 7.01 | 7.88 | 6.49 | 7.83 |
| S625 | 7.95 | 8.15 | 8.95 | 8.87 | 8.65 | 8.41 | 8.31 | 8.09 | 8.23 | 8.30 | 8.19 | 8.11 | 9.07 | 9.08 | 7.98 | 8.17 | 8.27 | 8.27 | 7.95 | 8.24 |
| S630 | ND | 6.37 | 8.33 | 7.38 | 6.37 | 6.92 | ND | ND | ND | 6.30 | ND | 6.49 | 8.17 | 7.30 | ND | 6.19 | ND | 6.42 | ND | ND |
| S633 | ND | 6.33 | ND | 6.57 | ND | 6.78 | ND | 6.16 | ND | 6.23 | ND | 6.34 | ND | 6.51 | ND | 6.32 | ND | 6.32 | ND | ND |
| S636 | ND | 6.57 | 6.45 | 7.98 | 6.59 | 8.13 | ND | 6.55 | ND | 6.47 | ND | 6.70 | 6.32 | 7.73 | ND | 6.65 | ND | 6.48 | ND | ND |
| S639 | ND | ND | 6.45 | 7.72 | 6.59 | 7.71 | ND | ND | ND | ND | ND | ND | 6.32 | 7.10 | ND | ND | ND | ND | ND | 6.05 |
| S642 | 6.32 | 6.35 | 7.27 | 7.78 | 7.32 | 7.89 | 6.50 | 6.22 | 6.39 | 6.31 | 6.22 | 6.18 | 6.87 | 7.21 | 6.27 | 6.40 | 6.33 | 6.36 | 6.63 | 6.60 |
| S643 | ND | 6.15 | 7.69 | 7.34 | 8.26 | 7.36 | ND | 6.13 | ND | 6.23 | ND | 6.07 | 7.15 | 7.03 | 6.20 | 6.11 | ND | 6.25 | ND | 6.05 |
| S652 | ND | ND | ND | ND | ND | 6.57 | ND | ND | ND | ND | ND | ND | ND | ND | ND | ND | ND | ND | ND | ND |
| S697 | ND | 6.16 | ND | 6.48 | ND | 6.93 | ND | ND | ND | ND | ND | ND | ND | ND | ND | ND | ND | ND | ND | ND |
| S700 | ND | 8.75 | ND | 9.17 | ND | 9.35 | ND | 8.38 | ND | 8.74 | ND | 8.50 | ND | 8.38 | ND | 8.38 | ND | 8.68 | ND | 8.90 |

Colour scale: ND (6.00) – 11.00

ND (6.00) 11.00

Suppl. Table 1. Mean phosphorylated peptide intensities (in log10 scale) obtained for individual sample from direct (pipeline #1) and enriched analysis processed (pipeline #2) are expressed in the shades of blue. ND corresponds to not detected signals or signals with intensity below $1\times10^6$.

**Supplementary Table2. Sequence coverage of DVL3**

| | Analysis | Replicate 1 | Replicate 2 | Replicate 3 | Sample | Experiment |
|---|---|---|---|---|---|---|
| Control 1 | Direct | 84.21 | 87.18 | 75.44 | 93.66 | 95.68 |
| | Enriched | 75.84 | 79.62 | 51.01 | | |
| CK1ε | Direct | 75.3 | 70.72 | 79.76 | 90.42 | |
| | Enriched | 68.83 | 61.13 | 69.5 | | |
| NEK2 | Direct | 85.83 | 76.25 | 82.32 | 94.06 | |
| | Enriched | 77.46 | 73.68 | 61.4 | | |
| PLK1 | Direct | 72.2 | 79.76 | 74.9 | 91.36 | |
| | Enriched | 70.58 | 62.08 | 57.35 | | |
| Aurora A | Direct | 78.54 | 76.79 | 83.27 | 93.12 | |
| | Enriched | 72.47 | 68.96 | 52.09 | | |
| Control 2 | Direct | 82.73 | 79.62 | 80.03 | 92.85 | |
| | Enriched | 67.07 | 69.77 | 59.92 | | |
| TTBK2 | Direct | 78.68 | 73.41 | 77.46 | 91.36 | |
| | Enriched | 76.11 | 66.4 | 65.86 | | |
| CK2α | Direct | 85.16 | 75.3 | 82.46 | 93.25 | |
| | Enriched | 74.09 | 68.96 | 65.32 | | |
| RIPK4 | Direct | 89.74 | 75.71 | 76.79 | 93.39 | |
| | Enriched | 72.33 | 65.59 | 52.5 | | |
| PKCδ | Direct | 73.82 | 78.41 | 83.4 | 91.23 | |
| | Enriched | 56.95 | 68.69 | 55.74 | | |

Sequence coverage of DVL3 obtained for individual kinases in each replicate, in individual sample (sum of all three replicates) and throughout the experiment.
